# piRNAs Coordinate poly(UG) Tailing to Prevent Aberrant and Permanent Gene Silencing

**DOI:** 10.1101/2021.01.30.428010

**Authors:** Aditi Shukla, Roberto Perales, Scott Kennedy

## Abstract

Noncoding RNAs have emerged as mediators of transgenerational epigenetic inheritance (TEI) in a number of organisms. A robust example of RNA-directed TEI is the inheritance of gene silencing states following RNA interference (RNAi) in the metazoan *C. elegans*. During RNAi inheritance, gene silencing is transmitted by a self-perpetuating cascade of siRNA-directed poly(UG) tailing of mRNA fragments (pUGylation), followed by siRNA synthesis from poly(UG)-tailed mRNA templates (termed pUG RNA/siRNA cycling). Despite the self-perpetuating nature of pUG RNA/siRNA cycling, RNAi inheritance is finite, suggesting that systems likely exist to prevent permanent RNAi-triggered gene silencing. Here we show that, in the absence of Piwi-interacting RNAs (piRNAs), an animal-specific class of small noncoding RNA, RNAi-based gene silencing can become essentially permanent, lasting at near 100% penetrance for more than five years and hundreds of generations. This permanent gene silencing is mediated by perpetual activation of the pUG RNA/siRNA TEI pathway. Further, we find that piRNAs coordinate endogenous RNAi pathways to prevent germline-expressed genes, which are not normally subjected to TEI, from entering a state of permanent and irreversible epigenetic silencing also mediated by perpetual activation of pUG RNA/siRNA cycling. Together, our results show that one function of *C. elegans* piRNAs is to insulate germline-expressed genes from aberrant and runaway inactivation by the pUG RNA/siRNA epigenetic inheritance system.

## Introduction

Small noncoding RNAs, such as PIWI-interacting RNAs (piRNAs), microRNAs (miRNAs), and small interfering RNAs (siRNAs) are key regulators of gene expression in eukaryotes. Small RNAs are bound by Argonaute proteins and, together, these ribonucleoprotein complexes target and silence complementary mRNAs^1,2^. piRNAs are an animal-specific class of genomically encoded and germline-expressed small noncoding RNA, which are bound by the PIWI clade of Argonaute proteins and are essential for germ cell function in many species of animals. One widely conserved function of piRNAs is to silence mobile genetic elements termed transposons^3,4^.

The functions of some piRNAs, however, remain enigmatic. For instance, most mammalian piRNAs do not exhibit complementarity to transposable elements^5–8^ and loss of Miwi, one of three PIWI proteins encoded by the mouse genome, results in male sterility, but does not cause transposon mobilization^9^. Thus mammalian piRNAs have important biological functions, in addition to transposon silencing. Similarly, the biological functions of most *C. elegans* piRNAs (also known as 21U-RNAs^10^) are mysterious. *C. elegans* piRNAs are bound and stabilized by PRG-1, one of two PIWI clade Argonaute proteins encoded by the *C. elegans* genome^11–16^. Surprisingly, fewer than 1% of transposon families are upregulated in a *prg-1* mutant^16^, which lacks all piRNAs^11–16^, indicating that transposon silencing is not the sole or possibly even major function of *C. elegans* piRNAs.

While *C. elegans* piRNAs do not play a vital role in transposon silencing, they do repress the expression of a number of germline-expressed mRNAs^11,13–17^, as well as germline-expressed transgenes^18–20^. Current models posit that PRG-1/piRNA complexes drive gene and transgene silencing by binding complementary mRNAs and recruiting RNA-dependent RNA Polymerases (RdRPs)^14,17^. RdRPs use target mRNAs as templates to synthesize 22G-siRNAs, which are antisense to mRNA templates, 22nt in length, and begin with a guanosine^21^. These piRNA-dependent 22G-siRNAs are bound by a worm-specific clade of Argonaute proteins (WAGOs) and, together, these ribonucleoprotein complexes mediate target mRNA silencing^21^. Interestingly, PRG-1 and its bound piRNAs interact with >16,000 mostly germline-expressed mRNAs^22,23^. However, fewer than 100 of these mRNAs undergo piRNA-dependent gene silencing^16^, indicating that PRG-1 and piRNAs do not silence most of the mRNAs to which they are bound. Indeed, several recent studies show that *C. elegans* piRNAs can promote the expression of some mRNAs by preventing aberrant siRNA-mediated gene silencing^15,16,24,25^. Why some genes become aberrantly silenced in a *prg-1* mutant and how PRG-1 and piRNAs normally prevent this silencing is a mystery.

PRG-1/piRNA-induced silencing of some loci, such as transgenes, can be heritable^18,19^. This heritable silencing is initiated by PRG-1 and piRNAs; however, once established, it can be maintained in the absence of PRG-1 and piRNAs for many generations^18–20^. Such heritable gene silencing, in the absence of initiating triggers, is an example of transgenerational epigenetic inheritance (TEI), which in *C. elegans* is also known as RNA-induced epigenetic silencing (RNAe)^18,20^. piRNA-induced transgenerational silencing correlates with both the heritable expression of histone 3 lysine 9 trimethylation (H3K9me3) at genomic loci undergoing RNAi inheritance and of 22G-siRNAs antisense to mRNAs transcribed from these loci. The process also depends on nuclear factors, such as the nuclear-localized WAGO HRDE-1, the HP1 homolog HPL-2, and histone methyltransferases^18,19^, suggesting that 22G-siRNAs and repressive post-translational histone modifications likely contribute to the heritable gene silencing initiated by piRNAs.

Double-stranded RNAs (dsRNAs) can also initiate TEI in *C. elegans*^18, 19,26–31^ via a conserved gene silencing program termed RNA interference (RNAi)^26^. In *C. elegans*, RNAi begins when dsRNA is processed into siRNAs, which are bound by the Argonaute RDE-1^32,33^. Together RDE-1/siRNA complexes bind target mRNAs via Watson-Crick base-pairing between the mRNA and the siRNA, resulting in target mRNA cleavage by the ribonuclease RDE-8^34^. RdRPs then use fragmented mRNAs to generate 22G-siRNAs, which, like piRNA-directed 22G-siRNAs, are bound by WAGOs to carry out gene silencing^21,35^. RNAi-directed silencing of germline-expressed mRNAs can be inherited (termed RNAi inheritance) and, like piRNA-directed heritable silencing of transgenes, RNAi inheritance depends on HRDE-1 and is correlated with the inheritance of 22G-siRNAs and repressive repressive histone modifications, such as H3K9me3^18,19,29^ and H3K27me3^36^. The convergence of RNAi-directed and piRNA-directed TEI on a common set of downstream gene silencing effector proteins, such as RdRPs and WAGOs (like HRDE-1), suggests that these two TEI pathways are related.

Maintenance of 22G-siRNA expression over generations after dsRNA-triggered TEI depends upon a recently discovered noncoding RNA modification^37^. mRNAs targeted by RNAi and cleaved by the RNAi machinery, such as the endonuclease RDE-8^34^, are modified with perfectly alternating 3’ uridine (U) and guanosine (G) repeats (termed poly(UG) or pUG tails) by the ribonucleotidyltransferase MUT-2/RDE-3^37,38^. pUG tails recruit RdRPs, which then use pUG RNAs as templates to generate 22G-siRNAs^37^. Generationally repeated rounds of pUG RNA-templated 22G-siRNA synthesis and 22G-siRNA-directed mRNA pUGylation (termed pUG RNA/siRNA cycling) appear to be the mechanism by which gene silencing memories are propagated across generations in *C. elegans*^37^. Although not yet tested, piRNA-directed TEI, which also requires RDE-3^18^, is also likely to depend upon pUG RNA/siRNA cycling. The self-perpetuating nature of the pUG RNA/siRNA cycling pathway hints that biological systems likely exist to limit and prevent this potentially dangerous pathway from mistargeting essential genes for generationally stable silencing. In support of this idea, RNAi inheritance is usually finite and recent studies have shown that the generational perdurance of RNAi inheritance is under genetic control^19,29,39–41^. For instance, the methyltransferase MET-2^40^ and the chromodomain protein HERI-1^41^ limit the number of generations that RNAi is inherited in *C. elegans*. Interestingly, HERI-1 is physically recruited to the chromatin of genes undergoing RNAi inheritance, suggesting that HERI-1 may play a direct role in limiting TEI^41^. Little else is known about how *C. elegans* might regulate and focus its potent, self-perpetuating, and potentially dangerous TEI pathways.

Here we show that piRNAs regulate the accuracy and duration of pUG RNA/siRNA-based transgenerational gene silencing in the *C. elegans* germline. In the absence of piRNAs, RNAi-initiated pUG RNA/siRNA cycling can become essentially permanent. Endogenous pUG RNA/siRNA cycling becomes disorganized, with genes normally expressed in the germline becoming inappropriate targets for poly(UG) tailing and undergoing gene silencing. We conclude that one function of *C. elegans* piRNAs is to protect germline-expressed genes from aberrant and runaway gene silencing by the pUG RNA/siRNA epigenetic inheritance pathway.

## Results

### Identification of mutations that cause RNAi to become essentially permanent

We previously conducted a forward genetic screen (Figure 1a) to identify factors that normally limit the duration of dsRNA-triggered TEI (RNAi inheritance) in *C. elegans*^41^. For this screen, we mutagenized animals harboring: (1) *oma-1*(*zu405*), a temperature-sensitive, gain-of-function mutation in the germline-expressed gene *oma-1* that causes embryonic arrest, unless *oma-1* is silenced by RNAi^42^; and (2) a *pie-1::gfp::h2b* transgene, which encodes a GFP::H2B fusion protein driven by a germline-expressed promoter (hereafter referred to as *gfp*). In wild-type animals, dsRNA-induced silencing of *oma-1* or *gfp* lasts for four to ten generations after initiating dsRNA triggers are removed^19,28,29,39,41^. Our screen identified 20 Heritable enhancers of RNAi (Heri) mutants that exhibited *oma-1* and *gfp* RNAi inheritance for seven or more generations longer than non-mutagenized controls^41^. *oma-1* and *gfp* expression was eventually restored within 20 generations after RNAi in 18 of our 20 mutants^41^. However, two mutants, *gg531* and *gg540*, were unique in that, more than five years, and hundreds of generations, after *gfp* and *oma-1* were initially targeted for silencing by RNAi, 100% of animals in these lineages continued to silence both *gfp* (Figure 1b) and *oma-1* (Figure 1c). Data presented below will show that the silencing of *gfp* and *oma-1* observed in *gg531* and *gg540* animals is epigenetic in nature. Henceforth, we refer to this remarkably stable epigenetic inheritance as “permanent silencing.” We conclude that systems exist to prevent dsRNA-initiated epigenetic inheritance from becoming essentially permanent.

**Figure 1.**
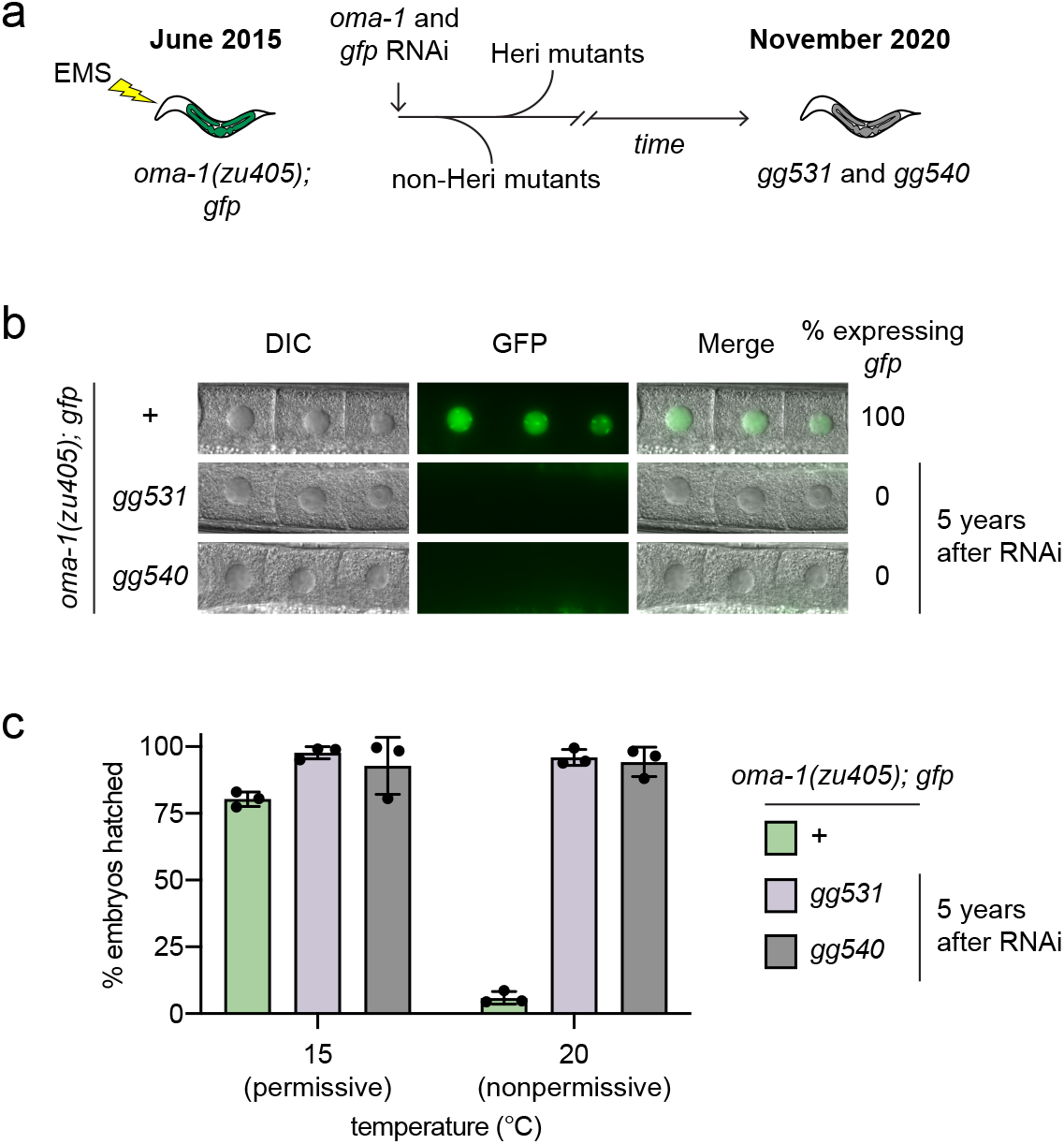
RNAi-mediated gene silencing can become permanent in *C. elegans*. **a,** *oma-1(zu405); gfp* animals were mutagenized with ethyl methanesulfonate (EMS) and fed bacteria expressing *oma-1* and *gfp* dsRNA (RNAi). In wild-type animals, *oma-1* and *gfp* RNAi can induce heritable silencing for several generations after dsRNA triggers are removed ^19,28,29,39,41^. *heritable enhancers of RNAi* (*heri*) mutations that extended both *oma-1* and *gfp* RNAi inheritance by seven or more generations were identified^41^. Whereas most Heri mutants re-expressed *oma-1* and *gfp* over the next 20 generations, *gg531* and *gg540* animals continue to inherit *oma-1* and *gfp* RNAi-mediated gene silencing at one hundred percent penetrance more than five years and hundreds of generations after RNAi. **b,** Fluorescence micrographs (63X oil objective) showing *gfp* expression in oocyte nuclei of animals of the indicated genotypes five years after *gg531* and *gg540* animals were exposed to *gfp* RNAi. >100 animals per genotype were scored using the 20X objective of a fluorescent microscope (Zeiss) and the % animals expressing *gfp* is indicated. If any GFP expression was detected, animals were scored as expressing *gfp*. **c,** *oma-1(zu405)* animals lay viable progeny at 15°C (permissive temperature), but lay arrested embryos at 20°C (nonpermissive temperature) unless *oma-1* is silenced by RNAi^42^. Therefore, the percentage (%) of *oma-1(zu405)* embryos that hatch [(# of hatched embryos / # embryos laid) x 100] at the nonpermissive temperature is a readout of *oma-1* silencing. For each replicate, % hatched embryos was measured and averaged for 6 individual animals of each of the indicated genotypes five years after *gg531* and *gg540* animals were exposed to *oma-1* RNAi. Error bars are standard deviation (s.d.) of the mean of the three biological replicates.

### PRG-1 limits RNAi inheritance

Whole-genome sequencing of *gg531* and *gg540* animals identified independent nonsense mutations in the gene *prg-1* (Figure 2a), which encodes one of two *C. elegans* PIWI clade Argonaute proteins^11–13^. Previous studies have shown that PRG-1 binds *C. elegans* piRNAs^11–13^ and, together, this complex can initiate transgenerational silencing of germline-expressed genes and transgenes via a mechanism involving piRNA-directed RdRP-based synthesis of 22G-siRNAs^18,19^. Our results suggest that, surprisingly, PRG-1 and piRNAs can also restrict transgenerational gene silencing initiated by exogenous dsRNA triggers. The following lines of evidence support this idea. First, we used genetic crosses to remove the silenced *gfp* allele from *prg-1(gg531)* animals and to reintroduce an expressed allele of *gfp*. We then tested these *gfp* expressing *prg-1(gg531)* animals for *gfp* RNAi inheritance and confirmed that these animals exhibited enhanced *gfp* RNAi inheritance (Figure S1a). Second, we asked whether animals harboring an independently isolated deletion allele, *tm872*, of *prg-1* exhibited an enhanced RNAi inheritance phenotype. *tm872* is a 640bp deletion (Figure 2a) in *prg-1* that removes most of the MID domain and part of the PIWI domain, two conserved domains found in Argonaute proteins^2^, and is, therefore, thought to represent a null allele of *prg-1*^13^. Indeed, the generational perdurance of RNAi-mediated gene silencing was dramatically enhanced in *prg-1(tm872)* animals (Figure 2b, Figure S1b). Together, these data confirm that loss of PRG-1 can enhance the generational perdurance of RNAi-induced gene silencing.

**Figure 2.**
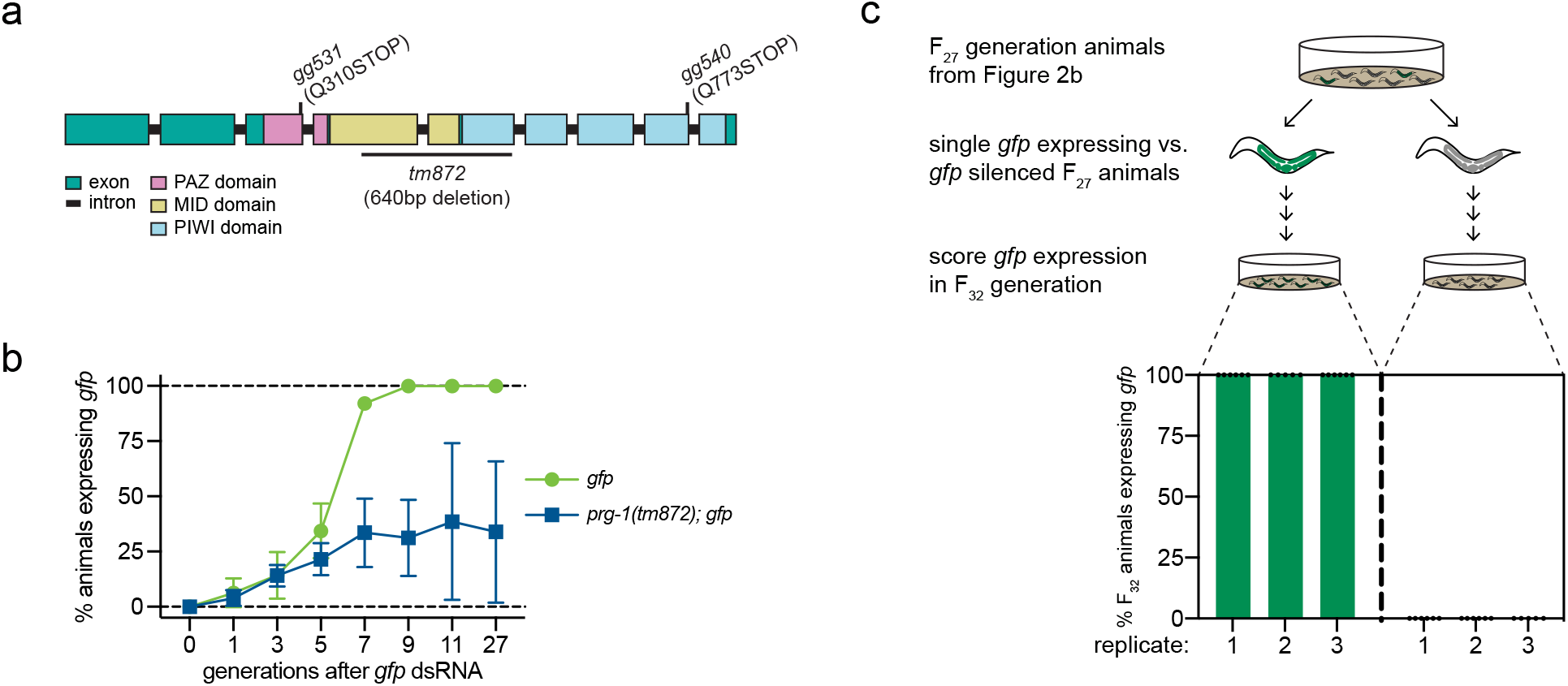
PRG-1 prevents permanent RNAi inheritance. **a,** Schematic of the *prg-1* gene, which encodes a PIWI clade Argonaute protein, indicating: (1) conserved PAZ, MID and PIWI domains; and (2) nature and locations of *prg-1* alleles used in this study*. prg-1(gg531)* and *prg-1(gg540)* are two nonsense alleles obtained from the forward genetic screen reported in Perales et al., 2018. *prg-1(tm872)* is an independently isolated partial deletion in *prg-1^13^. prg-1* coordinates were obtained from Wormbase release WS279. **b,** *gfp* RNAi inheritance assay performed on animals of the indicated genotypes. For each replicate, >50 animals per genotype were scored in each generation and the % of animals expressing *gfp* is shown. Error bars represent s.d. of the mean for three biological replicates. **c,** 5-6 *prg-1(tm872)* animals (represented by each point) in which *gfp* was either re-expressed or still silenced were singled 27 generations after *gfp* dsRNA treatment from each of the three biological replicates shown in panel **b**. Lineages were maintained for five additional generations and then *gfp* expression was scored.

Of note, some, but not all, *prg-1(tm872)* animals continued to inherit *gfp* silencing 27 generations after dsRNA treatment (Figure 2b). Given that the *prg-1* mutants we identified in our genetic screen continue to show 100% penetrant silencing five years after RNAi, we wondered whether, following *gfp* RNAi, a subset of *prg-1(tm872)* animals enters a state of permanent silencing in which they, and all of their progeny, exhibit fully penetrant gene silencing. To test this idea, we isolated individual *prg-1(tm872); gfp* animals in which *gfp* expression was either restored or still silenced 27 generations after *gfp* RNAi (Figure 2b), propagated lineages established from these individuals for an additional five generations and then scored *gfp* expression in each of the lineages (Figure 2c). This analysis showed that 27 generations after RNAi, individuals had entered one of two epigenetic states: either they and all of their progeny expressed *gfp* or they and all their progeny did not (Figure 2c, Figure S1c). For unknown reasons, the percentage of *prg-1* animals entering the permanently silenced state varied in different experiments (Figure 2b, Figure S1a, Figure S1b); however, in all cases, those animals that entered the permanently silent state never exited (Figure 2c, Figure S1c). We conclude that, in the absence of PRG-1, dsRNA triggers one of two epigenetic states: finite gene silencing, in which gene expression is eventually restored after several generations, or permanent gene silencing, in which targets of dsRNA are silenced forever.

We next asked if PRG-1 inhibits the initiation or maintenance of the RNAi-induced permanently silenced state. We crossed *prg-1(tm872)* animals in which *gfp* was silenced 13 generations after *gfp* RNAi to wild-type animals. We then monitored *gfp* expression in the *prg-1(+)* and *prg-1(tm872)* progeny of this cross, which were all homozygous for *gfp*. Only *prg-1(tm872)* progeny continued to silence *gfp* ~15 generations after outcross (Figure 3a), indicating that PRG-1 inhibits maintenance of permanent RNAi-induced gene silencing. To ask if PRG-1 inhibits the initiation of permanent silencing, we treated *gfp* expressing *prg-1(gg531/+)* heterozygous animals, which are wild-type for PRG-1 activity^19^, with *gfp* RNAi and scored the inheritance of *gfp* silencing in the *prg-1(+)* or *prg-1(gg531)* homozygous lineages of this cross. After ten generations, some *prg-1*(*gg531*), but no *prg-1*(+), lineages were still inheriting *gfp* silencing (Figure 3b), indicating that PRG-1 does not inhibit the initiation of permanent silencing, and confirming that PRG-1 inhibits the maintenance of *gfp* silencing after *gfp* RNAi. Taken together, these data show that PRG-1 acts in inheriting generations to prevent RNAi-initiated gene silencing from becoming permanent.

**Figure 3.**
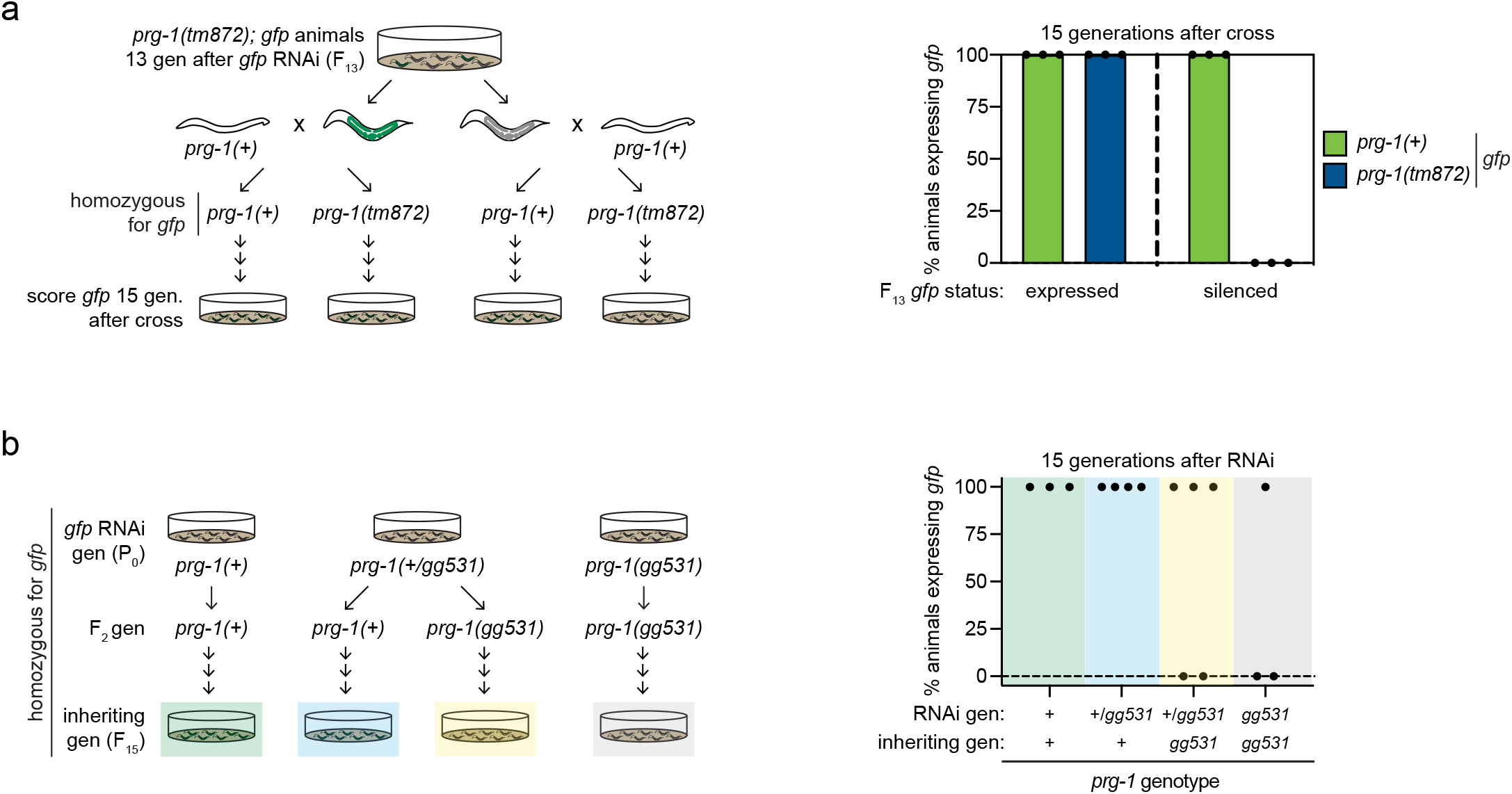
PRG-1 antagonizes the maintenance of heritable silencing. **a,** *prg-1(tm872); gfp* animals were fed *gfp* dsRNA. 13 generations (gen) after dsRNA treatment, three animals in which *gfp* was re-expressed and three animals in which *gfp* was still silenced were singled and independently crossed to *prg-1(+)* (e.g. wild-type) males. Lineages (represented by each point) that were homozygous for *gfp* were established from one *prg-1(+)* and one *prg(tm872)* F_2_ progeny from each of the crosses, maintained for 15 generations, and then scored for *gfp* expression. 30-80 animals were scored per lineage. **b,** *gfp* expressing animals of the indicated genotypes were exposed to *gfp* dsRNA (RNAi gen). Lineages (represented by each point) were established from singled F_2_ progeny of dsRNA-treated animals and *gfp* expression was scored in these lineages 15 generations after dsRNA treatment (inheriting gen). 50 animals were scored per lineage.

*C. elegans* PRG-1 possesses endonuclease (Slicer) activity *in vitro*^14^, which is dependent on an evolutionarily conserved DDH motif (catalytic triad)^2,14^, but whose biological function is not yet known. CRISPR/Cas9 mediated mutation of the catalytic triad, in a manner that disrupted PRG-1 Slicer activity *in vitro*^14^ (Figure S2a), did not result in permanent gene silencing after RNAi (Figure S2b). Thus, Slicer activity is not required for PRG-1 to limit RNAi inheritance and, therefore, the biological purpose of this activity remains enigmatic. Further, whereas our data show that PRG-1 suppresses permanent silencing triggered by exogenous dsRNA, the other *C. elegans* PIWI clade Argonaute, PRG-2, whose function is not yet known^11–13^, did not have a role in limiting RNAi inheritance (Figure S3).

### Permanently silenced alleles are paramutagenic

Paramutation is an epigenetic gene silencing phenomenon, first documented in plants^43^, whereby the epigenetic state of one allele is transmitted to another allele of the same gene. A related process occurs in the *C. elegans* germline, where it is also known as RNA-induced epigenetic silencing (RNAe)^18,20^. While conducting genetic crosses with *gg531* and *gg540* animals, we noticed that the permanently silenced *oma-1* and *gfp* alleles in *gg531* and *gg540* animals exhibited paramutagenic properties. For instance, when we crossed *prg-1(gg531)* animals harboring permanently silenced *gfp* or *oma-1* alleles to animals harboring expressed alleles of *gfp* or *oma-1* (identified by linked mutations in *dpy-10* or *dpy-20*, respectively), expressed alleles were converted to the silenced state (Figure 4a, Figure S4). This *trans*-silencing was highly penetrant, specifically when transmitted via the female germline (Figure S4), and was permanently maintained in lineages lacking PRG-1 (Figure 4a). We conclude that genes undergoing permanent silencing in *prg-1* mutant animals are paramutagenic, emphasizing the irreversible and permanent nature of RNAi silencing that can occur in the absence of PRG-1.

**Figure 4.**
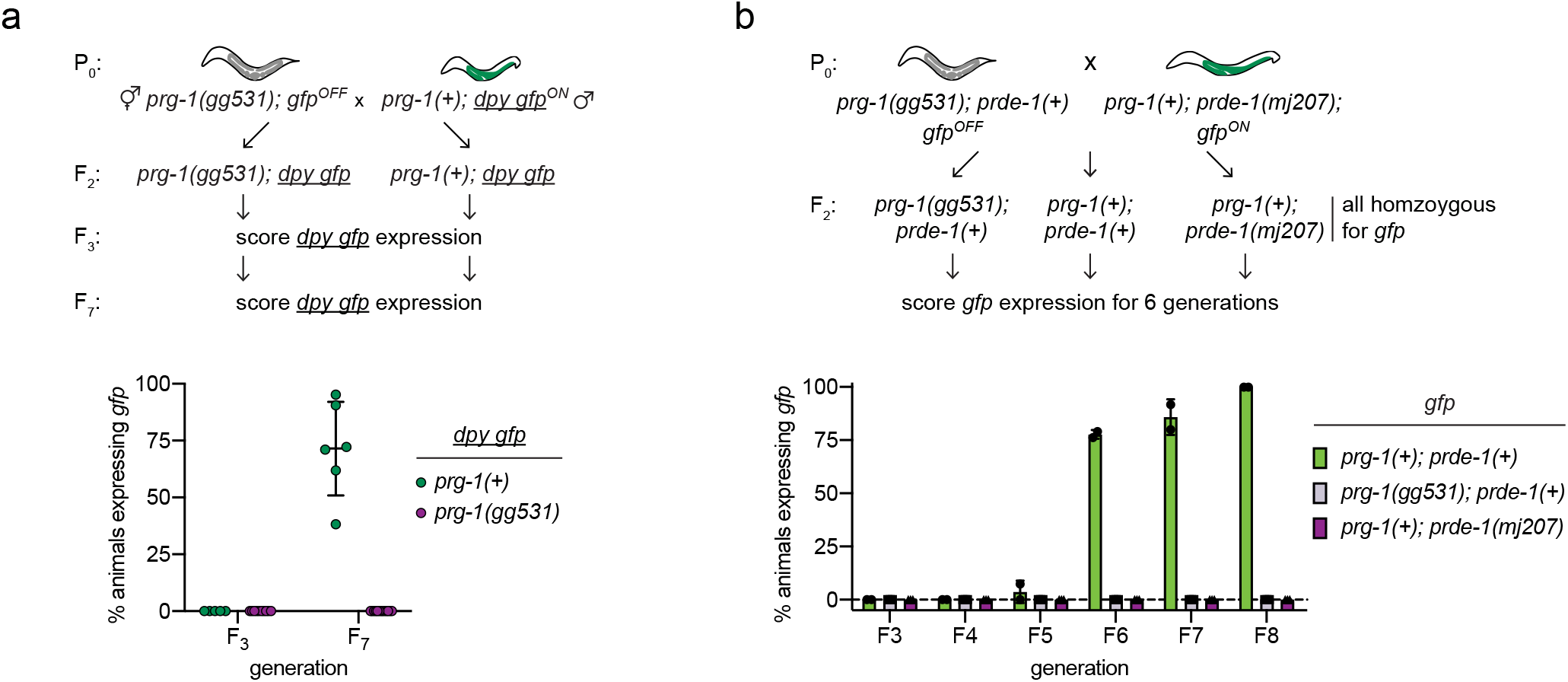
Permanently silenced alleles are paramutagenic. **a,** *prg-1(gg531)* hermaphrodites harboring a permanently silenced allele of *gfp (gfp^OFF^)* were crossed to *prg-1(+)* males harboring an expressed allele of *gfp* that was marked by a tightly linked *dpy-10(e128)* mutation (<u>*dpy gfp*</u>^*ON*^). Lineages (represented by each point) were established from singled F_2_ progeny of the indicated genotypes and *gfp* expression was scored in the indicated generations. 30-100 animals were scored per lineage in each generation. Error bars are s.d. of the mean. **b,** *prg-1(gg531); gfp^OFF^* hermaphrodites were crossed to *prde-1*(*mj207*); *gfp*^ON^ males. F_2_ animals were singled, genotyped for *prg-1(gg531)* and *prde-1(mj207)* and lineages of the indicated genotypes were established from these animals. *gfp* expression was scored after the indicated number of generations. 30-90 animals were scored per generation for each genotype. Error bars are s.d. of the mean.

### piRNAs limit RNAi inheritance

PRG-1 is a PIWI clade Argonaute protein that binds piRNAs in the *C. elegans* germline^11–13^. Therefore, the permanent silencing and paramutation we observe in animals lacking PRG-1 is likely due to the loss of piRNA function in the germline. In support of this idea, one of the twenty mutant strains identified by our genetic screen harbored a nonsense mutation, *gg530*, in the *prde-1* gene (Figure S5a), which encodes a nuclear-localized protein required for the production and/or stability of some piRNA precursor transcripts^44^. To further test the idea that PRDE-1, and, therefore, piRNAs prevent the maintenance of permanent silencing, we asked whether animals harboring *mj207*, an independently isolated null allele of *prde-1*^44^, also exhibited enhanced RNAi inheritance. We conducted *gfp* RNAi on *gfp* and *prde-1(mj207); gfp* animals and found that the loss of PRDE-1 had a complex and subtle effect on *gfp* RNAi inheritance (Figure S5b). Interestingly, 9 generations after dsRNA exposure, the loss of PRDE-1 caused a statistically significant enhancement of *gfp* RNAi inheritance (Figure S5b). Additionally, some *prde-1(mj207)* animals, but no control animals, continued to silence *gfp* 27 generations after RNAi (Figure S5b). Importantly, a related analysis that used paramutation to initiate gene silencing revealed a more dramatic and clearer role for PRDE-1 in the long-term maintenance of permanent silencing states. We crossed *prg-1(gg531)* animals harboring a permanently silenced *gfp* allele to *gfp* expressing *prde-1(mj207)* animals and isolated: (1) *prg-1(+); prde-1(+)*, (2) *prg-1(gg531)* or (3) *prde-1(mj207)* progeny from this cross, all of which were homozygous for *gfp* (Figure 4b). We then monitored *gfp* expression over generations in these lineages and, as expected based upon results described above (Figure 4a, Figure S4), paramutation-induced gene silencing was permanent in *prg-1(gg531)* but not in *prg-1(+); prde-1(+)* lineages (Figure 4b). *prde-1(mj207)* lineages behaved like *prg-1(gg531)* lineages in that they maintained *gfp* silencing for far more generations than *prg-1(+); prde-1(+)* lineages (Figure 4b). We conclude that PRDE-1, like PRG-1, limits the generational perdurance of RNAi inheritance, suggesting that piRNAs prevent permanent gene silencing in the *C. elegans* germline.

### Permanent silencing is driven by continuous siRNA production and continuous cTGS

Data in this section show that the permanent silencing of RNAi-targeted genes in animals lacking PRG-1 is due to the permanent activation of silencing pathways that are normally induced, but finite, after RNAi in wild-type animals. First, RNAi inheritance in wild-type animals is correlated with the heritable expression of 22G-siRNAs antisense to genes undergoing transgenerational silencing, which, in the case of *oma-1* RNAi inheritance, persist for 4-5 generations^19,29^. TaqMan-based siRNA quantification showed that *oma-1* (Figure 5a, Figure S6a) and *gfp* (Figure S6b) siRNAs were still expressed hundreds of generations after *gfp* and *oma-1* RNAi in both *prg-1(gg531)* and *prg-1(gg540)* animals. Second, during RNAi inheritance, genes are subjected to co-transcriptional gene silencing (cTGS), which, in the case of *oma-1* RNAi inheritance, persists for 4-5 generations^19,29^. qRT-PCR-based quantification of nascent *oma-1* pre-mRNA levels showed that *oma-1* remained co-transcriptionally silenced in *prg-1(gg531)* and *prg-1(gg540)* animals hundreds of generations after *oma-1* had been targeted for silencing by RNAi (Figure 5b). Third, three factors required for RNAi inheritance in wild-type animals – namely HRDE-1^18,19,29^, DEPS-1^45^, and ZNFX-1^45^ – were also needed for maintaining permanent silencing of *gfp* and *oma-1* in *prg-1(gg531)* and *prg-1(gg540)* animals (Figure 5c, Figure 5d, Figure S8). Together, these data show that the normally finite mechanism underlying RNAi inheritance in the *C. elegans* germline becomes permanent in the absence of PRG-1 and piRNAs.

**Figure 5.**
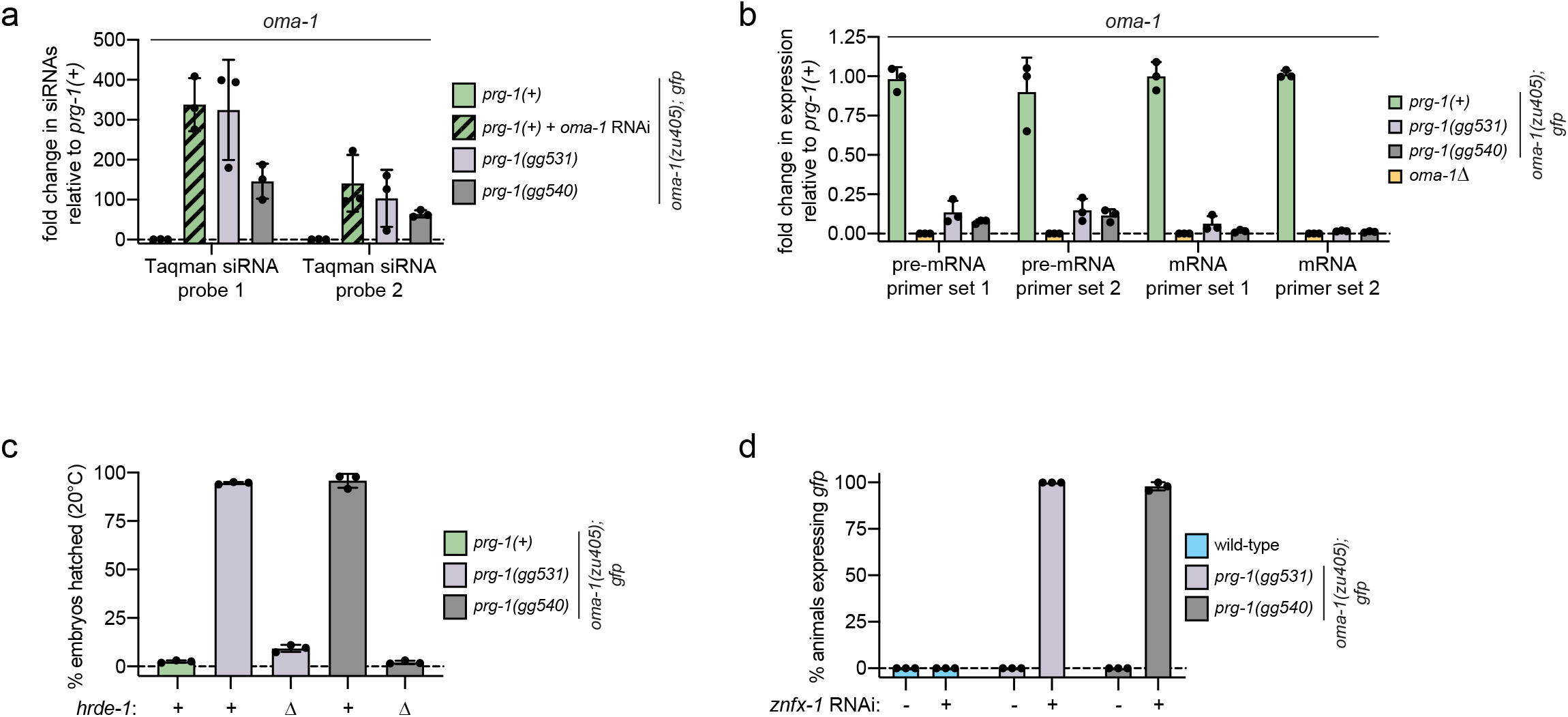
Permanent silencing is cotranscriptional and depends on known HRDE factors. **a,** Taqman-based qRT-PCR was used to quantify the expression of two different *oma-1* siRNAs (probes 1 and 2) in the following animals, all harboring *oma-1(zu405)* and *gfp*: (1) *prg-1(+)* animals +/- *oma-1* RNAi, (2) *prg-1(gg531)* animals, and (3) *prg-1(gg540)* animals, one year after *prg-1(gg531)* and *prg-1(gg540)* animals had been treated with *oma-1* RNAi. **b,** qRT-PCR using two primer sets (primer set 1 and 2) was used to quantify *oma-1* pre-mRNA and mRNA levels in animals of the indicated genotypes, two years after *prg-1(gg531)* and *prg-1(gg540)* animals had been treated with *oma-1* RNAi. Animals homozygous for *tm1396*, a 1515bp deletion in the *oma-1* gene (*oma-1Δ*), served as a negative control. *oma-1* pre-mRNA and mRNA levels were normalized to the germline-expressed gene *nos-3*. **c,** CRISPR/Cas9 was used to create a 2885bp deletion (Δ) that removes conserved domains in the *hrde-1* gene (Figure S7a) in *prg-1(gg531)* and *prg-1(gg540)* mutants. *oma-1* silencing was quantified as in Figure 1c for the indicated genotypes in three biological replicates. **d,** Animals of the indicated genotypes were fed bacteria expressing vector control (-) or dsRNA targeting the gene *znfx-1* (+) for two generations and *gfp* expression was then scored. Three biological replicates are shown, with 50 animals scored per treatment for each genotype. **a-d**, errors bars represent s.d. of the mean for three biological replicates.

### Perpetual pUG RNA/siRNA cycling causes permanent silencing

RDE-3 is a ribonucleotidyl-transferase required for RNAi^46,47^ and RNAi inheritance^37^ in *C. elegans*. RDE-3 adds poly(UG) or pUG tails to mRNAs targeted for silencing by RNAi (Figure 6a)^37^. These pUG tails recruit RdRPs, such as RRF-1, to pUG RNAs, which then serve as templates for antisense siRNA synthesis by RdRPs during RNAi and RNAi inheritance^37^. Generationally repeated sense/antisense cycles of pUG RNA-mediated siRNA biogenesis coupled with siRNA-directed mRNA pUGylation (pUG RNA/siRNA cycling) appear to mediate RNAi inheritance in the *C. elegans* germline (Figure 6a)^37^. However, despite the self-perpetuating nature of pUG RNA/siRNA cycling, the process typically only lasts a finite number of generations in wild-type animals^37^. The following data show that the permanent silencing of *oma-1* and *gfp* in *prg-1(gg531)* and *prg-1(gg540)* mutants is due to the perpetual activation of pUG RNA/siRNA cycling. First, we detected *oma-1* and *gfp* pUG RNAs in both *prg-1* mutant strains identified by our genetic screen, hundreds of generations after *oma-1* and *gfp* had been targeted for silencing by dsRNA (Figure 6b). These non-templated pUG tails were added to *oma-1* and *gfp* mRNA fragments (Figure S9a) at sites reminiscent of the pUGylation sites previously observed after RNAi in wild-type animals^37^. Second, pUG RNA expression correlated with permanent gene silencing. For instance, 3/3 *prg-1(tm872)* lineages that continued to silence *gfp* 33 generations after *gfp* RNAi (Figure 2c) expressed *gfp* pUG RNAs, while 3/3 lineages no longer silencing *gfp* did not (Figure 6c). Third, the poly(UG) polymerase RDE-3 was required for permanent silencing. A 484bp deletion introduced into the *rde-3* gene in *prg-1(gg531)* animals using CRISPR/Cas9 (Figure S7b) was sufficient to abrogate *gfp* and *oma-1* pUG RNA expression (Figure 6b), and to halt permanent *gfp* (Figure 6d) and *oma-1* silencing (Figure 6e). We conclude that permanent gene silencing in animals lacking piRNAs is driven by perpetual activation of the pUG RNA/siRNA pathway, indicating that PRG-1 and piRNAs normally limit the generational perdurance of pUG RNA/siRNA cycles.

**Figure 6.**
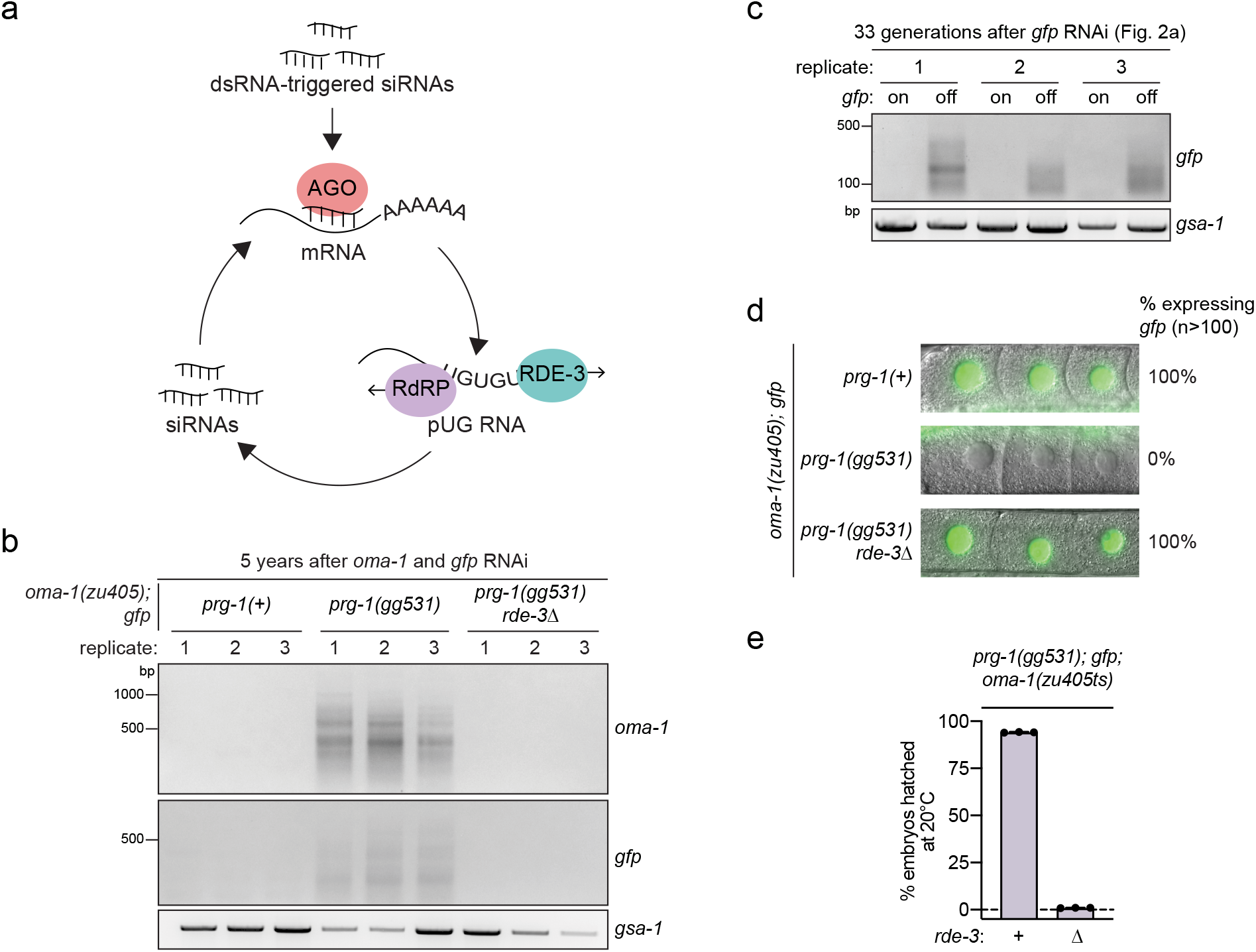
Permanent silencing is driven by perpetual pUG RNA/siRNA cycling. **a,** Model summarizing pUG RNA/siRNA cycling^37^. siRNAs derived from exogenous dsRNAs induce the cleavage of target mRNAs^34^. 5’ cleavage products are then modified with poly(UG) tails by the ribonucleotidyltransferase RDE-3. pUG RNAs serve as templates for 22G siRNA synthesis by RdRPs. A worm-specific clade of Argonaute proteins (termed WAGOs) binds 22G siRNAs and, together, this complex targets complementary mRNAs for: 1) transcriptional and translational silencing^29,35,60,61^ (not shown here), and 2) cleavage and subsequent *de novo* pUGylation. Cycles of pUG RNA-based siRNA production and siRNA-directed mRNA pUGylation maintain silencing over time and across generations. Despite the self-perpetuating nature of pUG RNA/siRNA cycling, for unknown reasons, cycling lasts for a finite number of generations in wild-type animals. **b,** Three biological replicates of pUG PCR (see Methods) to detect *oma-1* and *gfp* pUG RNAs were performed on total RNA isolated from animals of the indicated genotypes. A deletion (Δ) was introduced in the *rde-3* gene (Figure S7b) in *prg-1(gg531)* animals using CRISPR/Cas9. **b,** *gfp* pUG PCR was performed on total RNA isolated from *gfp* expressing and *gfp* silenced lineages that were established in Figure 2c. Animals were collected 33 generations after *gfp* dsRNA treatment. **b-c,** *gsa-1*, which has an 18nt genomically encoded pUG repeat in its 3’UTR, serves as a loading control. **d,** Fluorescence micrographs showing *gfp* expression in the oocytes of *oma-1(zu405); gfp* animals vs. *prg-1(gg531)* mutants with or without *rde-3* deletion (Figure S7b). >100 animals of each genotype were scored. **e**, *oma-1* silencing was measured in biological triplicate by quantifying the % embryos hatched at 20°C for animals of the indicated genotypes as described in Figure 1c.

### piRNAs prevent pUG RNA-based permanent silencing of germline-expressed genes

Recent studies have found that one function of *C. elegans* piRNAs is to coordinate endogenous 22G-siRNA systems in the germline^15,16,24,25^ and that, in the absence of this coordination, some germline-expressed genes, most strikingly the replication-dependent histones, undergo 22G-siRNA-dependent aberrant gene silencing^15,16^. We wondered if the mechanism underlying these previously documented cases of aberrant gene silencing in *prg-1* mutants^15,16^ might be aberrant pUG RNA/siRNA cycling. *his-10/14/26, his-11/15/44* and *his-12/16/43* are three sets of three nearly identical replication-dependent histones genes that are subjected to aberrant silencing in *prg-1* mutants^15,16^. We first confirmed that mRNAs encoded by these genes were aberrantly silenced in *prg-1(gg531)* animals (Figure 7a). We next asked if this aberrant silencing was associated with aberrant histone pUG RNA production. Indeed, we detected *his-10/14/26, his-11/15/44*, and *his-12/16/43* pUG RNAs in *prg-1(gg531)* animals, but not in *prg-1(+)* control animals (Figure 7b). Like pUG RNAs produced in response to dsRNA, histone pUG RNAs consisted of 5’ fragments of histone mRNAs modified with non-templated pUG tails (Figure S9b). Deletion of *rde-3* (Figure S7b) in *prg-1(gg531)* animals restored histone gene expression to near wild-type levels (Figure 7a) and abolished histone pUG RNA expression (Figure 7b). Expression profiling studies have identified a number of genes, in addition to the replication-dependent histones, whose expression is downregulated in *prg-1* mutants^15,16^. We wondered if aberrant silencing of these genes might also be explained by aberrant mRNA pUGylation. We tested this idea for the predicted protein-coding gene *r03d7.2*, which becomes downregulated in *prg-1* mutants (Figure 7c)^15,16^, and found that *r03d7.2* mRNAs were, indeed, pUGylated (Figure 7d) and silenced (Figure 7c) in an RDE-3-dependent manner in *prg-1(gg531)*, but not *prg-1(+)*, animals. We observed similar results for 2 out of 3 additional protein-coding genes previously reported to be downregulated in *prg-1* mutants^15,16^. We conclude that the downregulation of some predicted protein-coding mRNAs in *prg-1* mutants is mediated by aberrant and perpetual pUG RNA/siRNA cycling.

**Figure 7.**
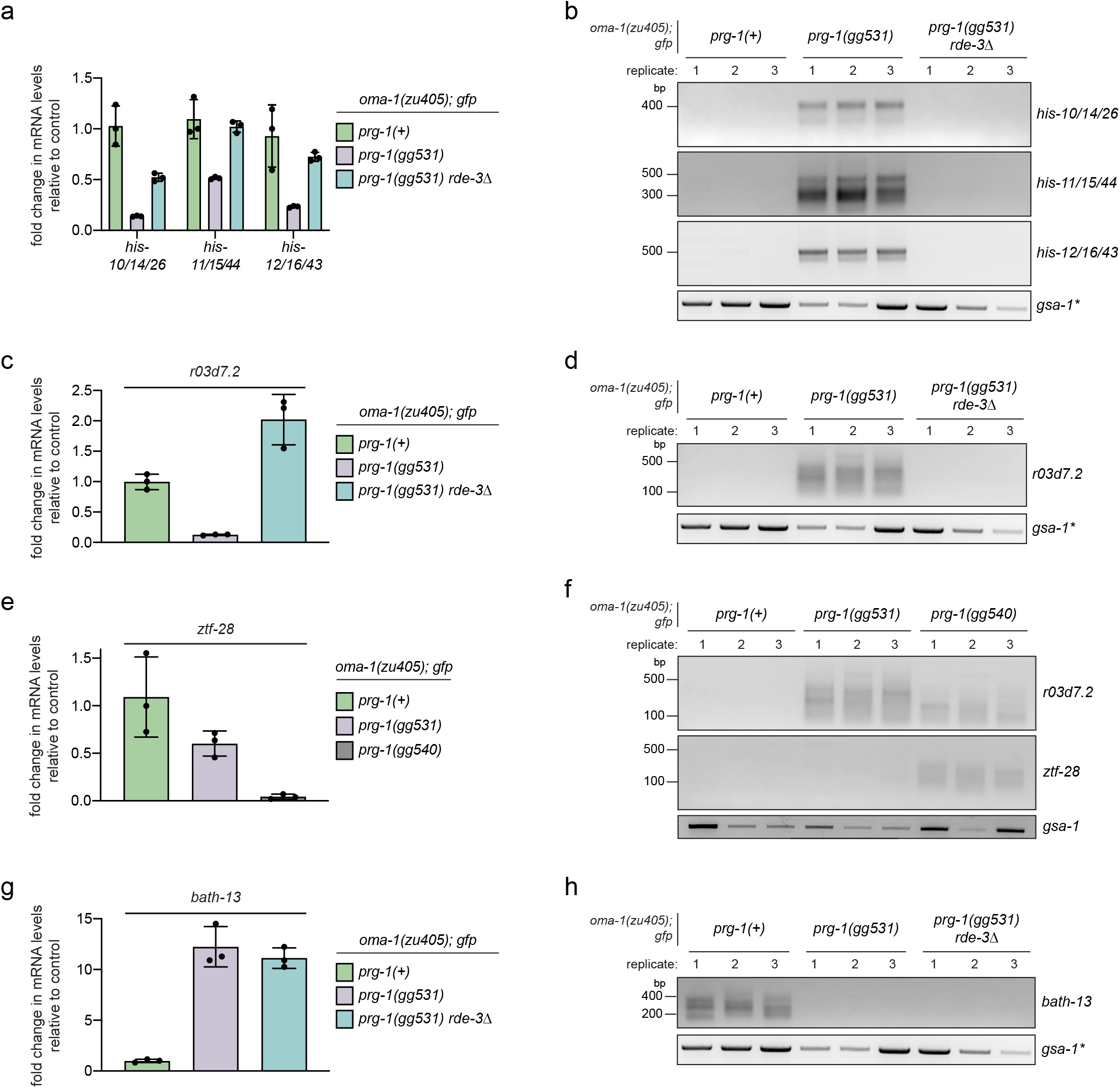
piRNAs coordinate the germline mRNA pUGylation system. **a, c, e, g,** qRT-PCR was performed in biological triplicate to quantify mRNA expression of the indicated genes using total RNA isolated from animals of the indicated genotypes. Results were normalized to the germline-expressed *nos-3* mRNA. Error bars are s.d. of the mean for three biological replicates. **b, d, f, h,** Gene-specific pUG PCR assays (see Methods) were performed in biological triplicate to detect pUGylation of the indicated mRNAs. *gsa-1* is a loading control. *The same RNA samples were used for panels b, d and h, so *gsa-1* data is identical in these panels.

Interestingly, while analyzing protein-coding genes for aberrant pUGylation, we found that mRNAs encoded by one gene, *ztf-28*, which were previously reported to be downregulated in *prg-1* mutants, were aberrantly silenced in *prg-1(gg540)* animals, but expressed at near wild-type levels in *prg-1(gg531)* animals (Figure 7e). Aberrant silencing of *ztf-28* in *prg-1(gg540)* animals correlated with *ztf-28* mRNA pUGylation in *prg-1(gg540)*, but not *prg-1(gg531)*, animals. Thus, while some genes, like the replication-dependent histone genes, are highly predisposed to aberrant pUGylation (and silencing) in the absence of piRNAs, other genes, like *ztf-28*, are subjected to aberrant pUGylation stochastically.

Finally, while some genes are known to undergo aberrant silencing in *prg-1* mutants, other genes are known to become upregulated^15,16^. We wondered if upregulation of these genes might be due to loss of mRNA pUGylation and, therefore, be indicative of global disorganization of the mRNA pUGylation system in the absence of piRNAs. We tested this idea for the *bath-13* gene, which has been shown to become upregulated in *prg-1* mutants^15,16^. We first confirmed that *bath-13* mRNA levels were upregulated in *prg-1(gg531)* mutants (Figure 7g). Additionally, we found that *bath-13* mRNA levels were similarly elevated in *prg-1(gg531)* and *prg-1(gg531); rde-3Δ* double mutant animals, suggesting that *prg-1* and *rde-3* normally act in the same genetic pathway to regulate *bath-13* (Figure 7g). We next asked if *bath-13* mRNA upregulation in *prg-1* mutants correlated with a loss of *bath-13* mRNA pUGylation. Indeed, we found that *bath-13* mRNAs are normally pUGylated in *prg-1(+)* animals and that this pUGylation was lost in *prg-1(gg531)* animals (Figure 7h). Taken together, the data suggest that PRG-1 and piRNAs coordinate gene expression programs in the *C. elegans* germline by focusing the activity of the poly(UG) polymerase RDE-3 activity to the appropriate mRNAs.

## Discussion

Here we show that in the absence of PRG-1 and piRNAs, dsRNA-triggered TEI, which normally lasts for only a finite number of generations, can become essentially permanent. This permanent silencing, whose fidelity of transmission approaches that of DNA-based inheritance, is driven by perpetual pUG RNA/siRNA cycling. Further, we find that, in the absence of piRNAs, the endogenous mRNA pUGylation system becomes disorganized, with some natural targets of RDE-3 escaping modification, and other mRNAs, including the replication-dependent histone RNAs, becoming novel substrates for RDE-3 (Figure S10a). Together, these results show that piRNAs prevent germline-expressed mRNAs from entering self-perpetuating pUG RNA/siRNA cycles, thus protecting these mRNAs from undergoing permanent and runaway transgenerational silencing.

How PRG-1/piRNAs regulate the targets of pUGylation in *C. elegans* remains a mystery, but here we propose some models that might explain this novel function of PRG-1/piRNAs. P granules are biomolecular condensates that form in *C. elegans* germ cells^48,49^ and promote germ cell fate specification by transmitting maternal mRNAs and epigenetic factors to the germline precursor cell during embryonic development^50^. The PRG-1/piRNA complex binds thousands of germline-expressed transcripts, and many of these interactions likely occur in P granules, where PRG-1 localizes^11,13^. Recent work showed that binding by PRG-1/piRNAs may serve to sequester mRNAs to P granules, preventing these mRNAs from interacting with silencing machinery in the cytoplasm^51^. Indeed, our work suggests this sequestration may prevent some transcripts from interacting with and, thus, being pUGylated by, RDE-3, which localizes to *Mutator* foci, distinct cytoplasmic perinuclear germline condensates required for 22G-siRNA amplification^47^. Interestingly, a related model has been proposed for piRNA function in *Drosophila* where piRNAs promote germline specification by binding to and, thereby, entrapping maternal mRNAs inside germ granules, which are biomolecular condensates constituting part of the fly germplasm^52^. Thus, the promotion of germline gene expression programs via sequestration of mRNAs within biomolecular condensates may be a conserved and ancient function of the animal piRNAs. Alternatively, PRG-1/piRNAs may coordinate the pUGylation system by competing with other RNAi pathways, including the exogenous RNAi pathway, for the same downstream silencing factors (RdRPs and WAGOs)^19,35,53,54^. Therefore, in the absence of piRNAs, components of this shared downstream silencing machinery are freed up to maintain aberrant pUG RNA/siRNA cycles. Related competition models have been proposed to explain why mutations in the ERI/DICER complex enhance exo-RNAi^55–57^, why endo-RNAi mutants show enhanced miRNA silencing^58^, and why animals lacking the MET-2 methyltransferase show prolonged RNAi inheritance^40^. Of note, it is also possible that a complex interplay between mRNA sequestration, competition between RNAi pathways and other PRG-1/piRNA-dependent phenomenon combine to explain how PRG-1/piRNAs coordinate mRNA pUGylation in *C. elegans*.

Why some mRNAs, but not others, become novel targets of pUGylation in the absence of piRNAs is also not known. Whereas exogenous dsRNA is likely the trigger that induces permanent silencing of mRNAs targeted by RNAi when PRG-1/piRNAs are absent, the molecular triggers initiating aberrant silencing of germline-expressed transcripts, such as the histone mRNAs, appear to be more complex. It was previously proposed that the lack of a poly(A) tail, which is characteristic of eukaryotic replication-dependent histone mRNAs, might predispose histone mRNAs to aberrant silencing in piRNA mutants^15^. However, we find that mRNAs that are thought to be polyadenylated, such as *r03d7.2*, can also be subjected to irreversible pUGylation and silencing (Figure 7d) indicating that the molecular signals triggering aberrant silencing in the absence of piRNAs remain to be identified. Our observation that genes differ in their intrinsic susceptibility to runaway silencing only adds an additional layer of complexity to the molecular basis of aberrant silencing. For instance, the *ztf-28* mRNA, which was previously reported to be aberrantly silenced in piRNA mutants^15,16^, was subjected to aberrant pUG RNA/siRNA-based silencing in just one of the two *prg-1* mutant strains identified from our forward genetic screen. Similarly, we find that 27 generations after experimental RNAi, some *prg-1(-)* lineages show permanent silencing of *gfp*, while others do not (Figure 2b). Together, these results hint at a molecular threshold that must be reached for an mRNA to become a target of pUG RNA/siRNA cycling and, thus, to become irreversibly silenced. According to this model, some mRNAs, like the histone mRNAs, are more highly predisposed to reaching this threshold and, therefore, runaway histone silencing is inevitable and highly penetrant in populations of *prg-1* mutants. Other mRNAs, like *ztf-28*, are less likely to reach this threshold and, therefore, runaway silencing of these mRNAs is less penetrant and stochastic. However, while genes may differ in their propensity to enter a state of permanent silencing, because pUG RNA/siRNA cycling is self-perpetuating, once this “event horizon” threshold is reached, gene silencing is complete and irreversible in all cases. One question that arises from these findings is whether expressed mRNAs ever spontaneously enter pUG RNA/siRNA cycling in wild-type animals, thereby creating epialleles. Given the variable and heritable nature of epigenetic states that we observe in the absence of piRNAs, it is enticing to consider if such epialleles could provide variation that allows *C. elegans* populations to adapt to changing environments. In support of this idea, spontaneous siRNA-maintained epimutations arise at a rate 25x higher than DNA mutations in *C. elegans* populations^59^.

The germline pUG RNA/siRNA gene silencing pathway is self-perpetuating and, therefore, potentially dangerous. If, for example, any germline mRNA was to mistakenly enter the pathway, it might never exit. Therefore, dedicated systems, such as PRG-1 and piRNAs, exist to limit and/or regulate pUG RNA/siRNA cycling. *prg-1* mutants exhibit fertility defects, including reduced brood size, that are exacerbated at elevated temperatures^11–13^. The aberrant silencing of essential genes that occurs in the absence of piRNAs has been linked to this sterility^24,25^, and genetic perturbations that restore expression to aberrantly silenced genes in *prg-1* mutants partially rescue the sterility defects of *prg-1* mutants^16^. These results, combined with our data showing that aberrant silencing in *prg-1* mutants is driven by pUG RNA/siRNA cycling, suggests that some of the germline defects associated with *prg-1* are due to disorganization of RDE-3-dependent pUGylation, indicating that coordination of pUG RNA/siRNA cycling is an important mechanism by which piRNAs promote germ cell function.

## Acknowledgements

We would like to thank past and present members of the Kennedy lab for helpful discussions of the data, and Marv Wickens for comments on the manuscript. Some strains were provided by the *Caenorhabditis* Genetics Center (CGC), which is funded by the NIH Office of Research Infrastructure Programs (P40 OD010440). Some strains were provided by the Mitani laboratory through the National BioResource Project (Tokyo, Japan), which is part of the International *C. elegans* Gene Knockout Consortium. A.S. (DGE1144152, DGE1745303) was supported by a NSF Graduate Research Fellowship. R.P. was supported by a Ruth L. Kirschstein National Research Service Award (1F32GM120919-01A1).

## Author contributions

A.S. collected all of the data presented in the manuscript. R.P. performed the forward genetic screen that identified the mutants characterized in this manuscript. A.S. and S.K. wrote the manuscript.

## Supplemental Figures Legends

**Figure S1.**
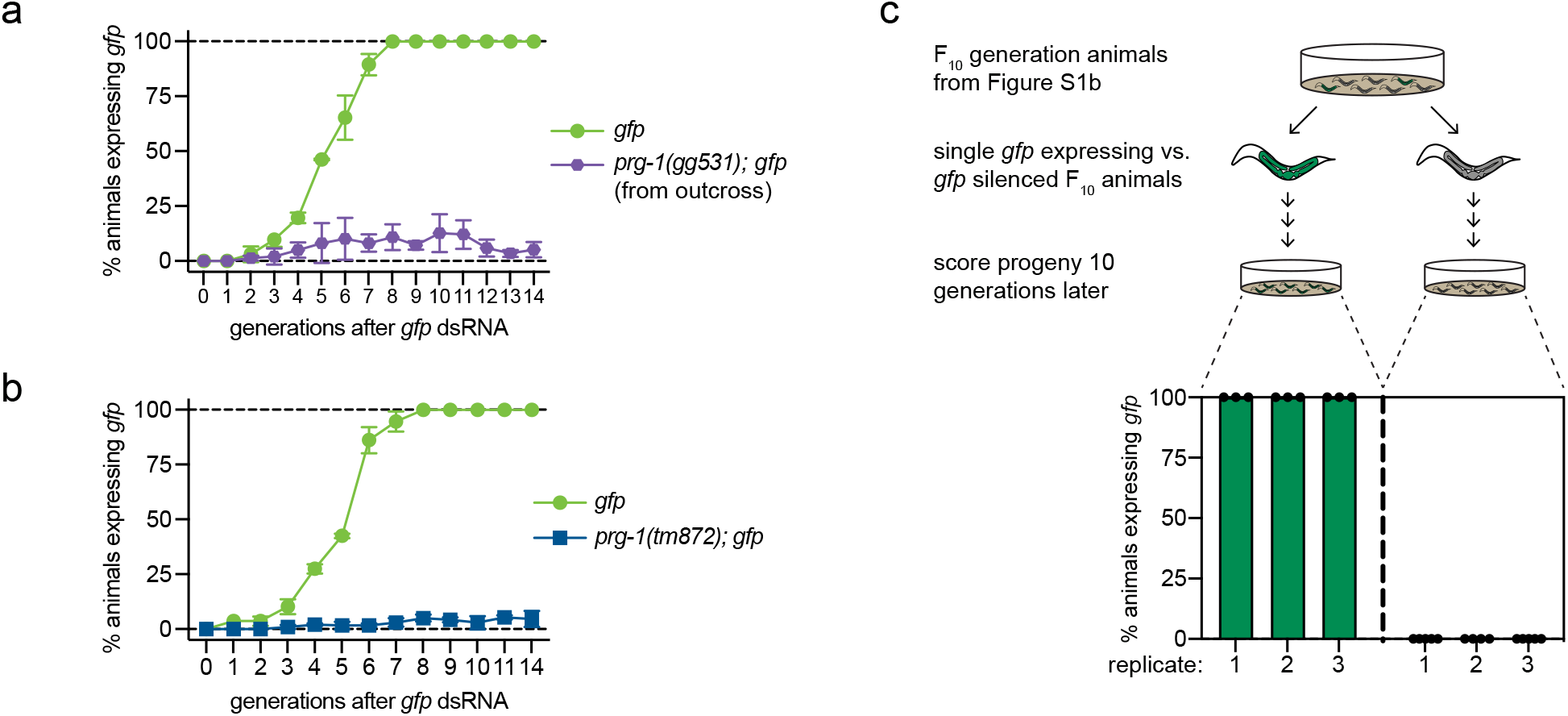
PRG-1 limits the generational perdurance of *gfp* RNAi inheritance. **a-b,** *gfp* RNAi inheritance was performed on animals of the indicated genotypes. Animals were fed *gfp* dsRNA and the percentage of animals expressing *gfp* was scored at the indicated number of generations after dsRNA treatment. Error bars represent s.d. of the mean of three biological replicates, with >50 animals counted per replicate for each genotype every generation. **a,** *prg-1(gg531)* mutant animals were outcrossed to remove the silent *gfp* allele and reintroduce an expressed *gfp* allele. **b,** This inheritance assay was performed two years before the inheritance assay shown in Figure 2b and employed a *prg-1(tm872)* strain that was nearing 100% sterility, so it was outcrossed with *gfp* expressing animals to generate the strain used for Figure 2b. **c,** 3-5 *prg-1(tm872)* animals (represented by each point) that re-expressed or continued to silence *gfp* were singled 10 generations after *gfp* dsRNA treatment from each biological replicate in the experiment shown in Figure S1b. Lineages established from these animals were maintained for 10 additional generations and then *gfp* expression was scored.

**Figure S2.**
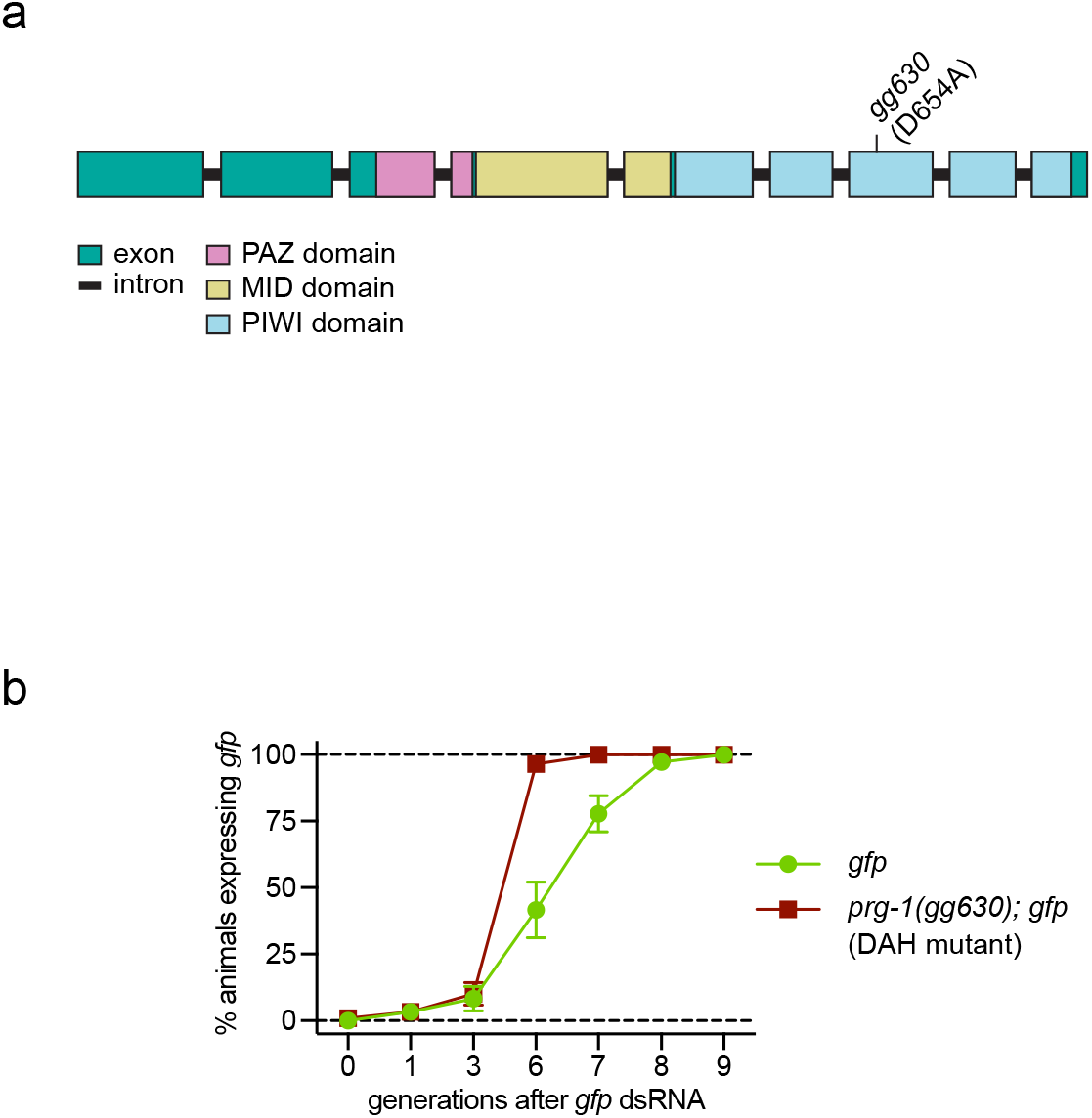
PRG-1 prevents permanent RNAi inheritance independent of its Slicer activity. **a,** An evolutionarily conserved aspartic acid-aspartic acid-histidine (DDH) motif (catalytic triad) mediates the Slicer activity of Argonaute proteins^2,14^. CRISPR/Cas9 was used to create a missense mutation (*gg630*) in *prg-1* that altered the second D (amino acid 654) in this catalytic triad to A, a mutation previously shown to abolish PRG-1 Slicer activity *in vitro*^14^. **b**, *gfp* RNAi inheritance assay performed on *gfp* expressing control animals vs. *prg-1(gg630)* (e.g. DAH) mutants. >50 animals were counted each generation for each genotype. Error bars represent s.d. of the mean of three independent biological replicates.

**Figure S3.**
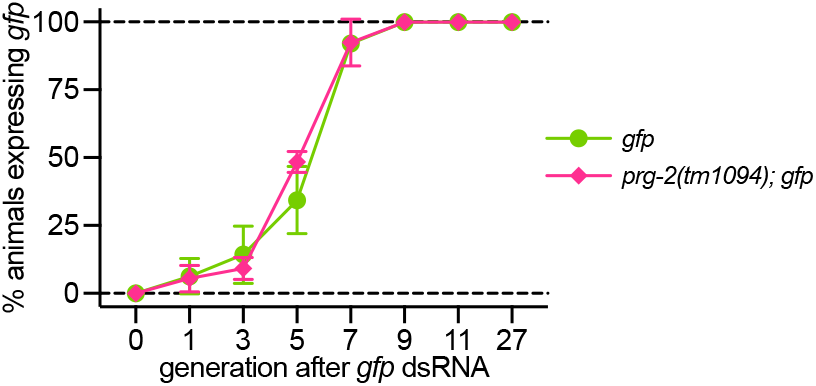
PRG-2 does not limit RNAi inheritance. PRG-2 is the second PIWI clade Argonaute encoded by the *C. elegans* genome, but does not seem to bind *C. elegans* piRNAs ^11–13^. Three biological replicates of a *gfp* RNAi inheritance assay were performed on *gfp* expressing control vs. *prg-2(tm1094); gfp* animals. >50 animals were counted each generation for each genotype. Error bars represent s.d. of the mean of three independent biological replicates.

**Figure S4.**
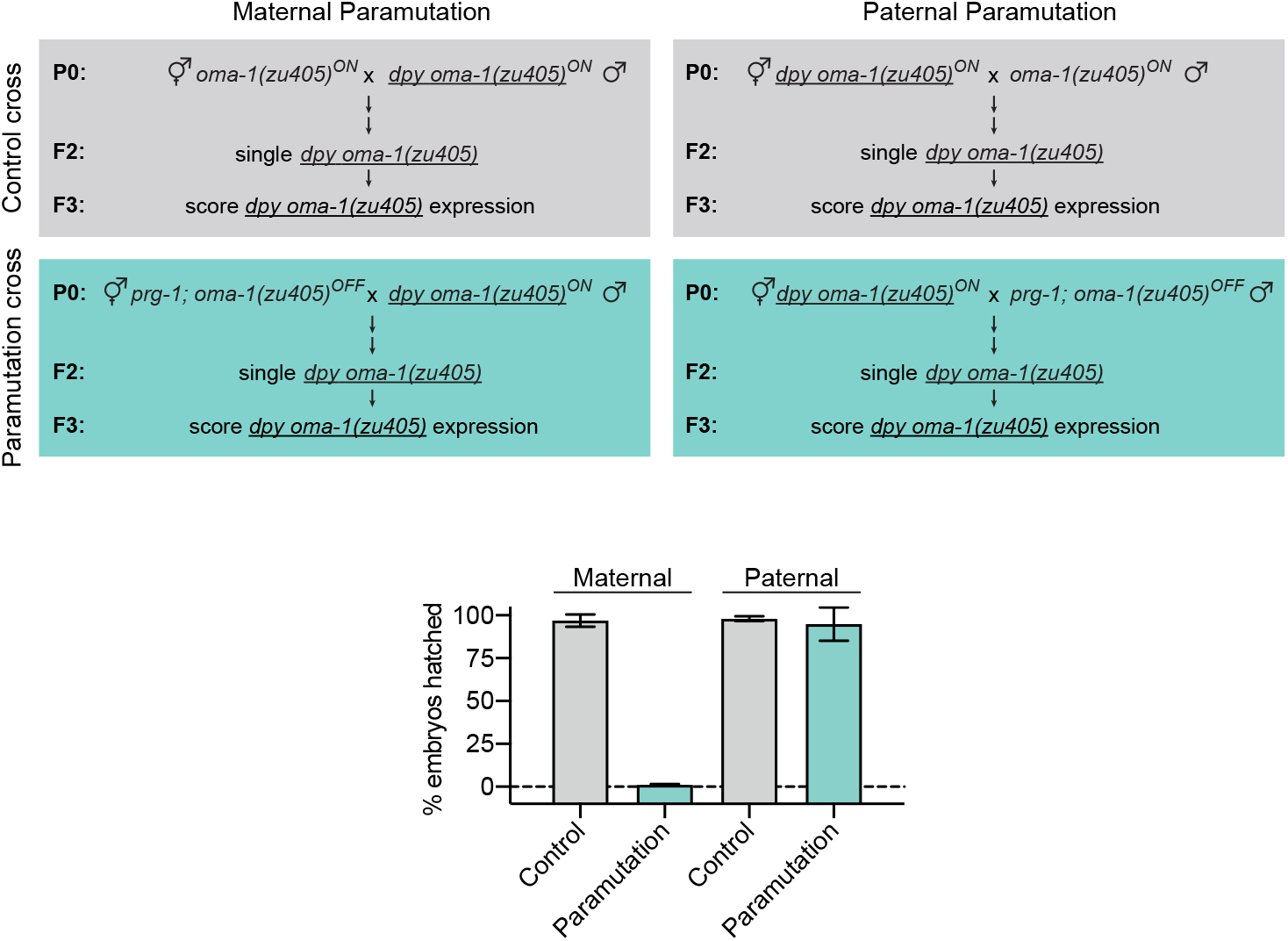
Permanently silenced *oma-1* alleles are paramutagenic and paramutation is more potent via the female germline. Paramutation cross: A permanently silenced allele of *oma-1(zu405) (oma-1(zu405)^OFF^)* was crossed into animals that harbored an expressed *oma-1* allele marked by a linked *dpy-20(e1282)* allele (<u>*dpy oma-1(zu405)*</u>^*ON*^). Permanently silenced alleles were introduced via the female germline (maternal paramutation) or the male germline (paternal paramutation). F_2_ animals that were homozygous for <u>*dpy oma-1(zu405)*</u> were singled and % hatched embryos were scored in the F_3_ generation. Control cross: <u>*oma-1(zu405)^ON^*</u> was crossed into *dpy oma-1(zu405)^ON^* animals via the male or female germline. For all crosses, four pairs of adult animals were mated, 20 <u>*dpy oma-1(zu405)*</u> F_2_ animals were isolated (from 5 different F_1_ animals), and % embryos hatched was scored in pools of 5 F_3_ animals derived from each F_2_ animal. % embryos hatched value is an average of all lineages derived from the same P_0_. Error bars represent s.d. of the mean.

**Figure S5.**
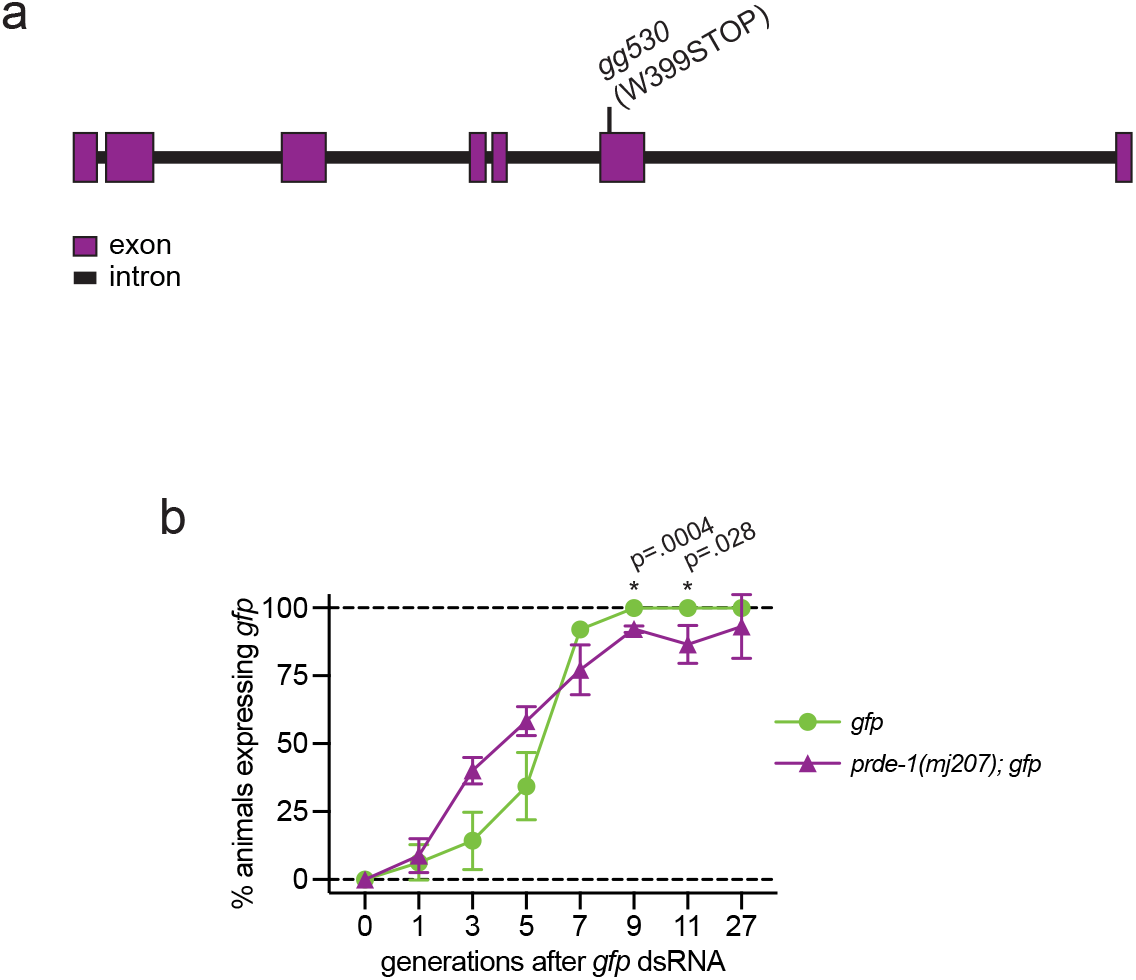
PRDE-1 may also prevent permanent RNAi inheritance. **a,** *gg530* was identified from the Heri screen described previously^41^ as showing enhanced inheritance of *oma-1* and *gfp* RNAi compared to unmutagenized animals. *gg530* mutants harbor the indicated nonsense mutation in the *prde-1* gene. **b,** *gfp* RNAi inheritance assay was performed on *gfp* expressing control vs. *prde-1(mj207)* animals. >50 animals were counted each generation for each genotype. Error bars represent s.d. of the mean of three independent biological replicates. Of note, *prde-1(mj207)* animals were tested for *gfp* RNAi inheritance alongside the *prg-1(tm872)* animals shown in Figure 2b. Therefore, the control data in these two figures is the same.

**Figure S6.**
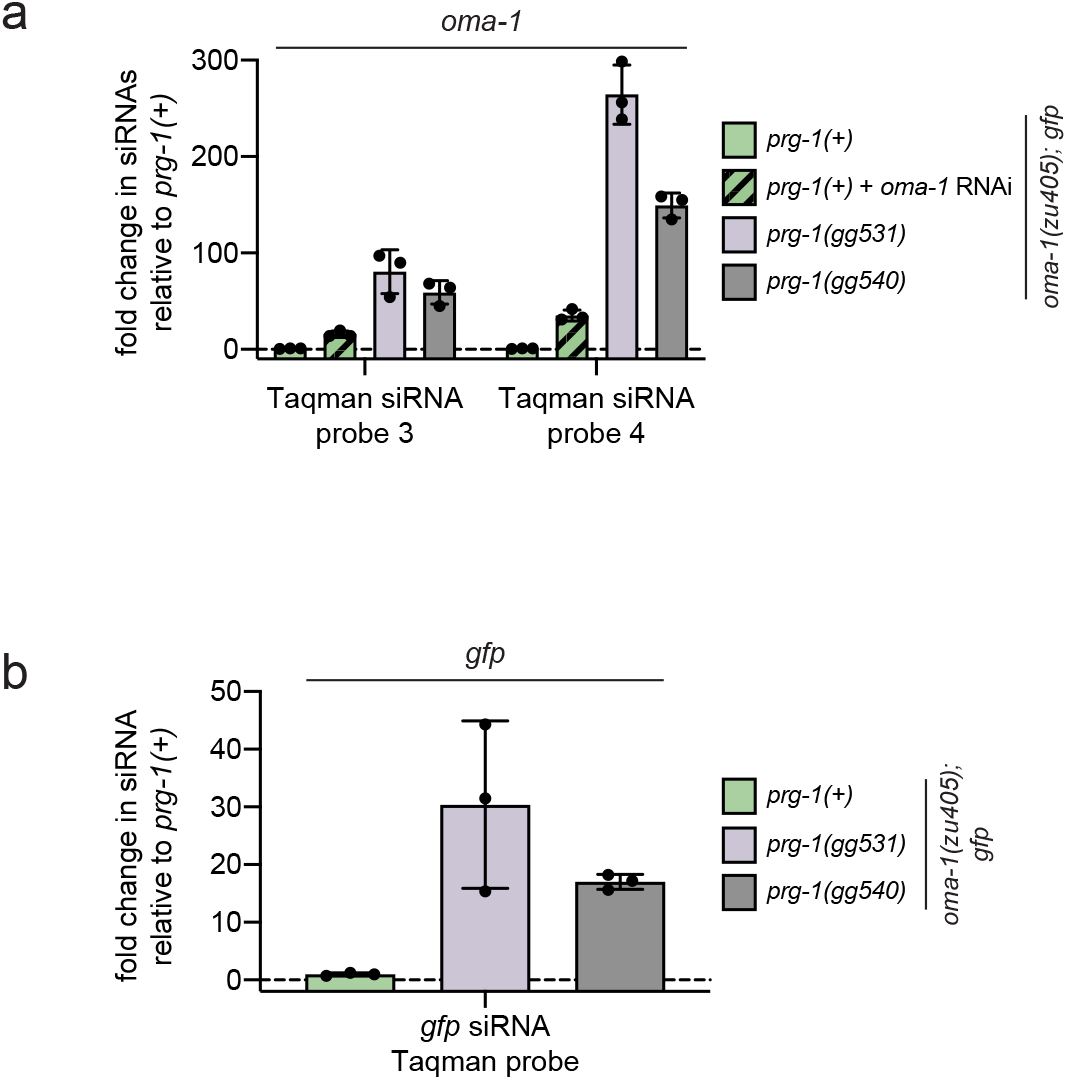
*prg-1(gg531)* and *prg-1(gg540)* mutants show elevated *oma-1* and *gfp* siRNA levels. Taqman-based qRT-PCR was used to quantify the expression of: **a,** two additional (see Figure 5a) *oma-1* siRNAs (probes 3 and 4) in the following animals, all harboring *oma-1(zu405)* and *gfp*: (1) *prg-1(+)* animals +/- *oma-1* RNAi, (2) *prg-1(gg531)* animals, and (3) *prg-1(gg540)* animals, one year after *prg-1(gg531)* and *prg-1(gg540)* animals had been treated with *oma-1* RNAi. **b,** a *gfp* siRNA in *prg-1(+), prg-1(gg531)* and *prg-1(gg540)* animals, one year after *prg-1(gg531)* and *prg-1(gg540)* animals had been treated with *gfp* RNAi.

**Figure S7.**
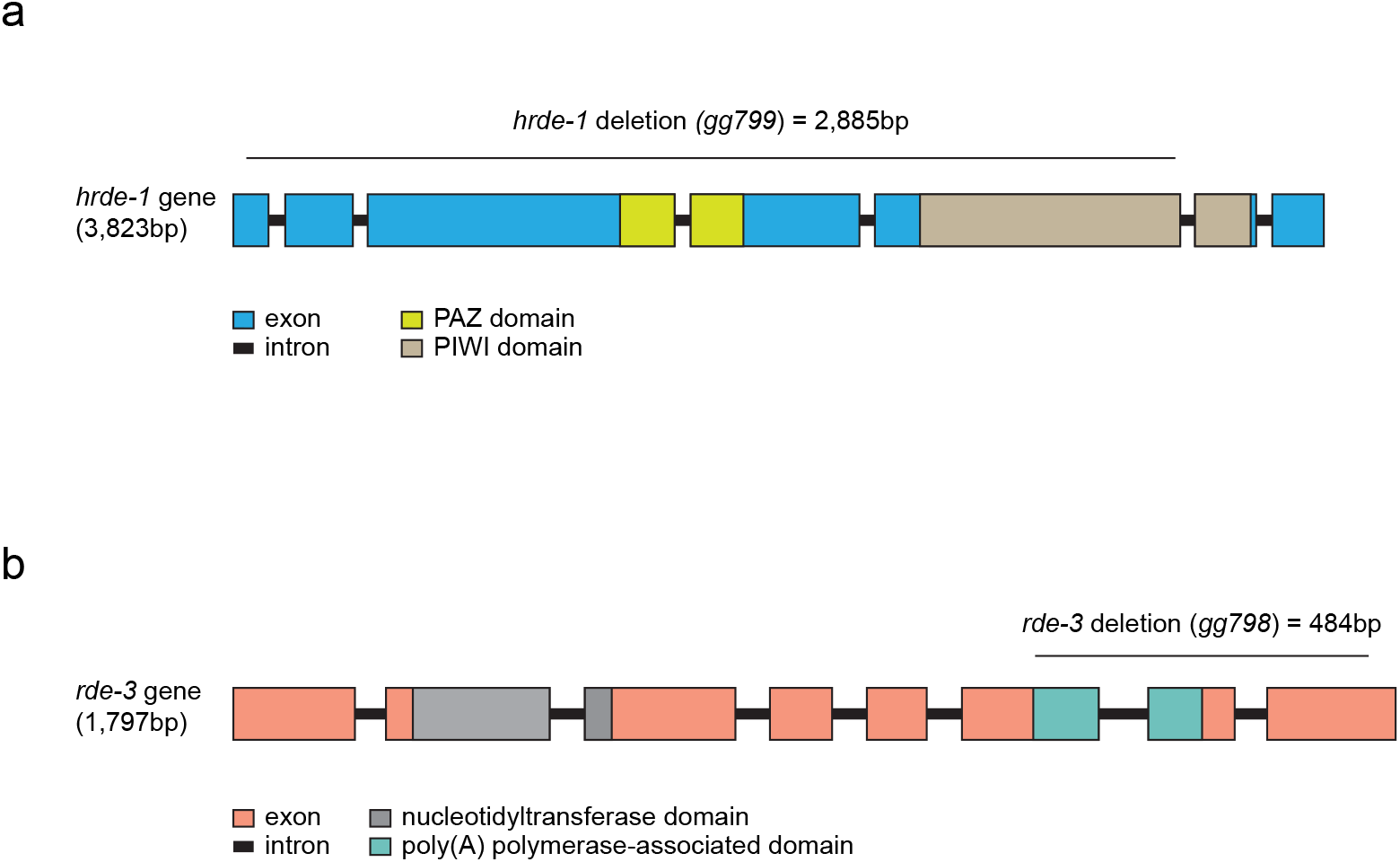
CRISPR/Cas9-induced deletions in *hrde-1* and *rde-3* genes. **a,** Schematic of the *hrde-1* gene, annotated with *gg800*, a 2885 deletion introduced into *prg-1(gg531)* and *prg-1(gg540)* mutants (Figure 5c). **b,** Schematic of the *rde-3* gene, annotated with *gg799*, a 484bp deletion introduced into *prg-1(gg531)* mutants using CRISPR/Cas9. This deletion introduces an out-of-frame mutation that results in a premature nonsense mutation in *rde-3*. For **a** and **b**, gene features and coordinates are from Wormbase release WS279.

**Figure S8.**
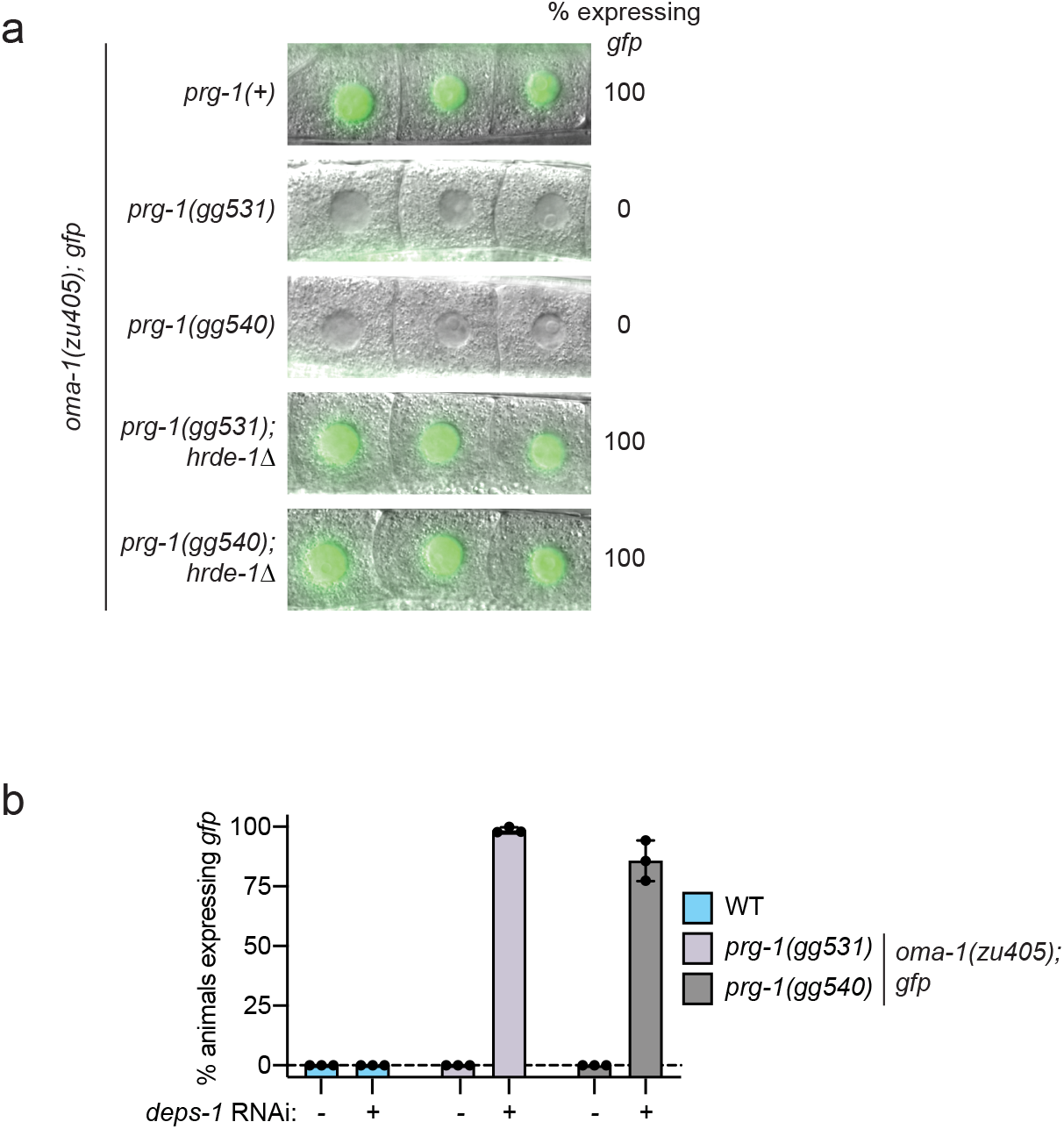
HRDE-1 and DEPS-1 are required for permanent RNAi inheritance in *prg-1(gg531)* and *prg-1(gg540)* mutants. **a,** *prg-1(gg531)* and *prg-1(gg540)* mutants with and without the *hrde-1* deletion (Δ) shown in Figure S7a were scored for *gfp* expression, alongside *oma-1(zu405); gfp* control animals. >100 animals were scored for each genotype. **b,** Control (*oma-1(zu405); gfp*) animals and *prg-1(gg531)* and *prg-1(gg540)* mutants were fed bacteria expressing empty vector control or dsRNA targeting the gene *deps-1* for two generations and then *gfp* expression was scored. Error bars represent s.d. of the mean of three biological replicates.

**Figure S9.**
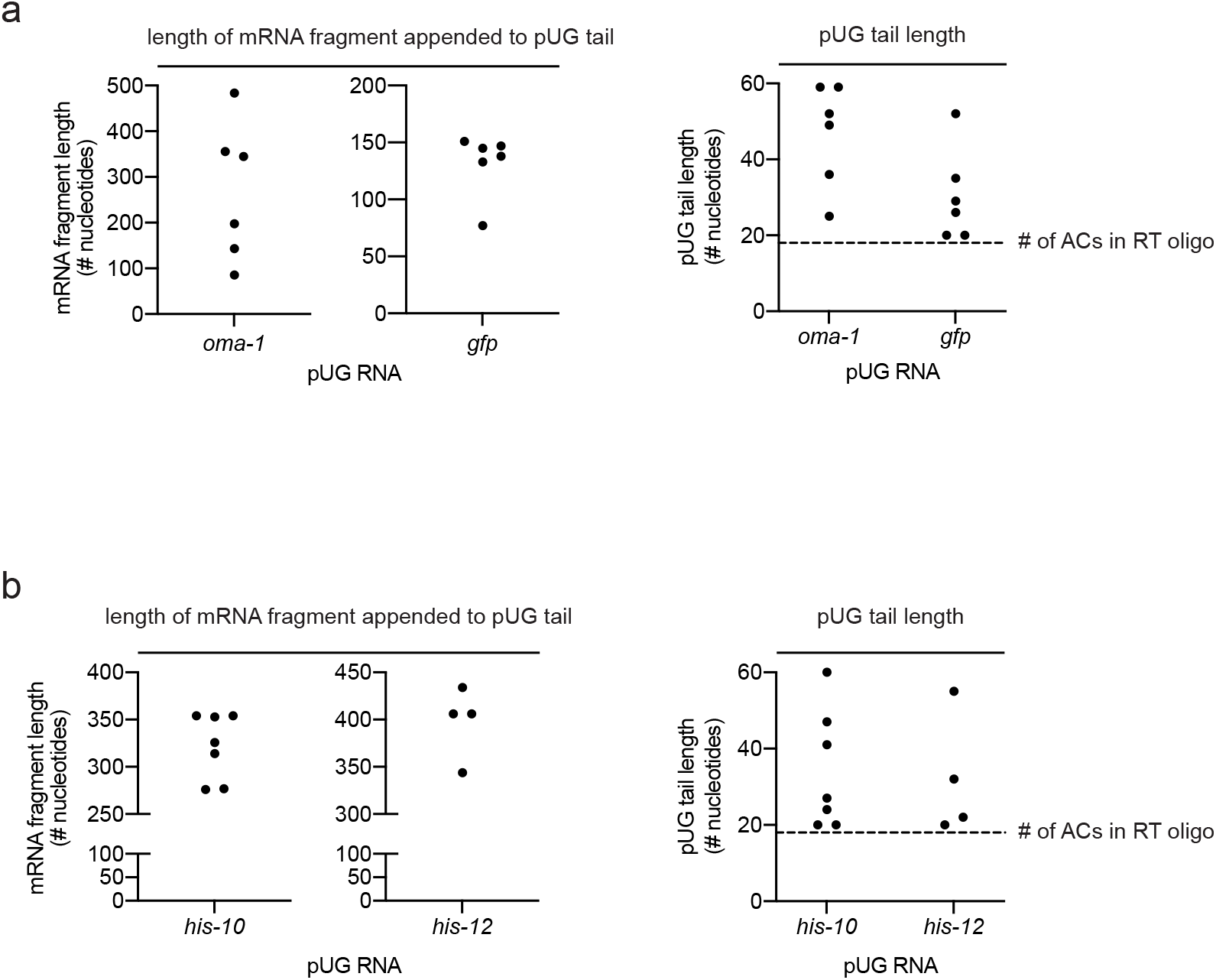
*oma-1, gfp* and histone pUG RNAs in *prg-1(gg531)* mutants. Sanger sequencing (see Methods) was performed on **a,** *oma-1* and *gfp* pUG RNAs; **b,** histone pUG RNAs detected in *prg-1(gg531)* mutants (Figures 6a and 7b, respectively). Sequencing showed that pUG PCR products in Figures 6a and 7b were spliced mRNA fragments modified with long 3’ nontemplated pUG tails. The length of spliced mRNA fragment modified with pUG tails and the lengths of pUG tails detected are shown. Only pUG RNAs harboring >18nt long pUG tails (i.e. the length of the RT oligo used to detect them) were analyzed.

**Figure S10.**
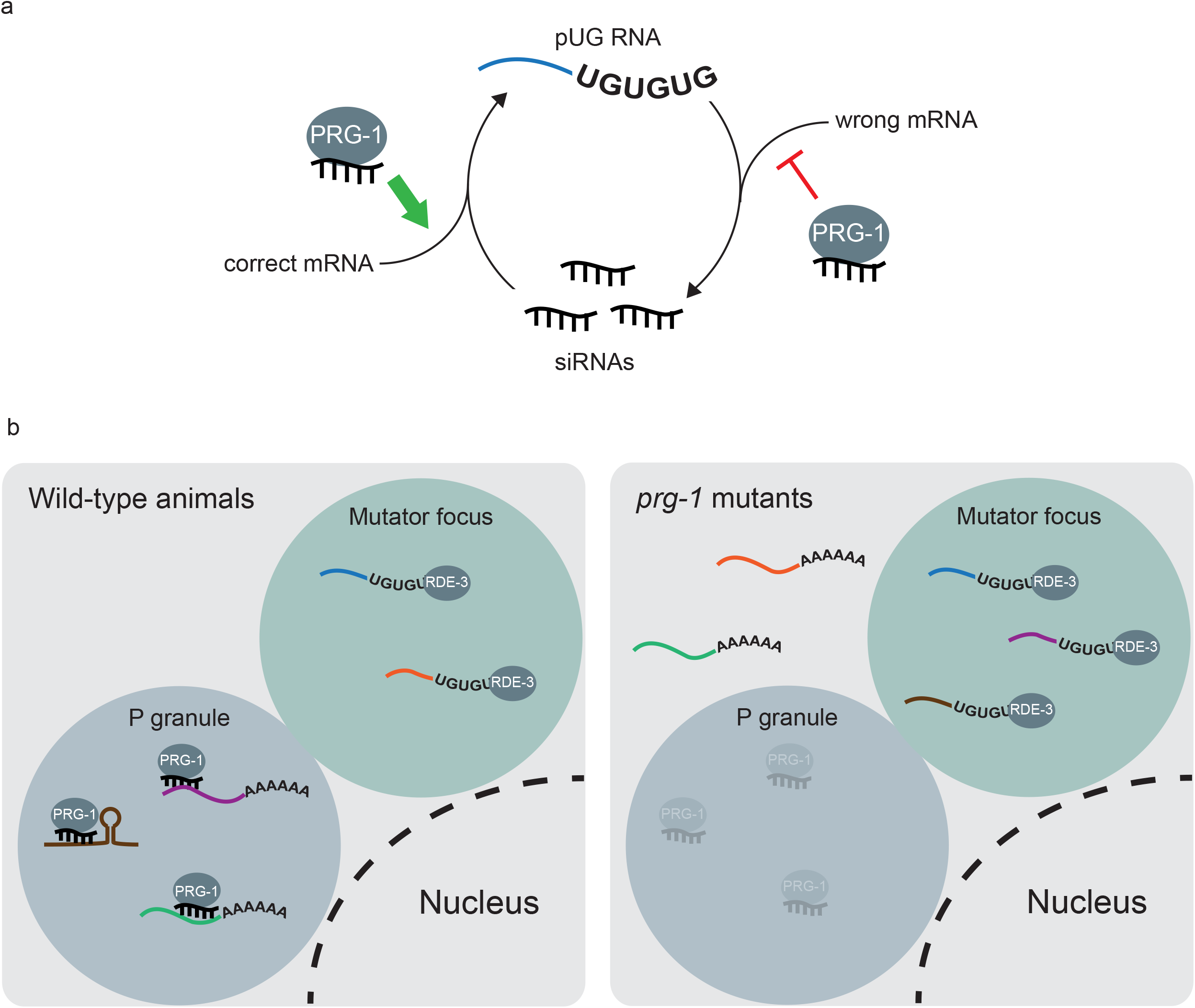
piRNAs coordinate pUG RNA/siRNA cycling. **a,** Our data suggests that piRNAs regulate germline gene expression by by: (1) protecting some mRNAs from pUGylation and, therefore, promoting their expression; and (2) by promoting pUGylation and, therefore, silencing of other mRNAs. Molecular signals dictating why some mRNAs are repressed by piRNAs while other mRNAs are protected, and the reason why the loss of piRNAs differentially affects the pUGylation status of these mRNAs, are not yet known. **b,** One way in which PRG-1/piRNAs may regulate the targets of pUGylation is by binding and sequestering some mRNAs within germline biomolecular condensates called P granules, where PRG-1 is known to localize^11,13^. mRNAs sequestered in P granules become inaccessible to the poly(UG) polymerase RDE-3, which localizes to distinct germline biomolecular condensates called *Mutator* foci^47^. According to this model, in the absence of piRNAs, some mRNAs escape P granules and enter *Mutator* foci, where they are cleaved by RDE-8 and pUGylated by RDE-3. The presence of aberrant mRNAs within *Mutator* foci disrupts pUGylation of other mRNAs by RDE-3.

## Methods

### Genetics

*C. elegans* culture and genetics were performed as described previously^1^. Strains were maintained at 20°C on Nematode Growth Medium (NGM) plates and fed OP50 *E. coli* bacteria, unless otherwise noted. Of note, animals expressing *zu405*, a gain-of-function, temperature-sensitive mutation in the gene *oma-1*^2^ were maintained at 15°C. Strains used in this study are listed in Table 1.

### RNAi and RNAi inheritance assays

To perform RNAi experiments, embryos were isolated via hypochlorite treatment (egg prep) of gravid adult hermaphrodites and dropped onto RNAi plates (standard NGM plates with 1 mM Isopropyl β-D-1-thiogalactopyranoside and 25ug/ml carbenicillin) seeded with HT115 *E. coli* bacteria expressing either L4440 (Addgene, #1654) empty vector control or L4440 carrying inserts to trigger the production of dsRNA (RNAi) against a gene of interest. The *oma-1* RNAi clone came from the *C. elegans* RNAi collection (Ahringer lab). The *gfp* RNAi clone was obtained from the Fire lab. The *znfx-1* RNAi clone was a custom clone made for this study.

To measure *oma-1(zu405ts)* silencing, embryos were isolated via hypochlorite treatment of gravid adult hermaphrodites and dropped onto *oma-1* RNAi plates. Animals were grown at 20°C for 2-3 days. 6 larval stage 4 (L4) or adult animals were singled for each strain/genotype and allowed to lay embryos overnight at 20°C. Adults were removed from plates on the next day and the total number of embryos laid was counted. On the following day, the total number of embryos that hatched was counted. % embryos hatched was then calculated as the (# of hatched embryos / # embryos laid) x 100. For *oma-1* RNAi inheritance experiments, the first generation of animals were treated with *oma-1* RNAi and then each generation, some adults were scored for *oma-1(zu405ts)* silencing while the remaining population was egg prepped and embryos were dropped onto NGM plates seeded with OP50 *E. coli* bacteria. This process was repeated every generation for the indicated number of generations.

For *gfp* RNAi and RNAi inheritance assays, embryos were dropped onto RNAi plates seeded with bacteria expressing either the L4440 empty vector control or *gfp* dsRNA. Once animals were gravid adults, they were washed off of plates using M9 + Triton X-100 buffer. 50-150 animals were placed onto a microscope slide and *gfp* expression was scored using the Plan-Apochromat 20 × /0.8 M27 objective on an Axio Observer.Z1 fluorescent microscope (Zeiss). Images were taken with the Plan-Apochromat 63 × /1.4 Oil DIC M27 objective. All image processing was done using Fiji ^3^. For *gfp* RNAi inheritance experiments, each generation, 50-150 gravid adults were scored for *gfp* expression and the remaining population of gravid adults were egg prepped and embryos were dropped onto NGM plates seeded with OP50 *E. coli* bacteria. This process was repeated every generation for the indicated number of generations.

### Paramutation crosses

Expressed *oma-1(zu405ts)* and *gfp* alleles were marked with tightly-linked *dpy-20(e1282)* and *dpy-10(e128)*, respectively, mutations to allow for differentiation of expressed/naive alleles from silent *oma-1(zu405ts)* and *gfp* alleles in *prg-1(gg531)* and *prg-1(gg540)* mutants. More specifically, *oma-1(zu405ts)* and *gfp* animals were crossed with *dpy-20(e1282)* and *dpy-10(e128)* animals, respectively, and 200-300 F_2_s were singled and genotyped to identify rare double mutants resulting from a crossover. Paramutation crosses for *oma-1(zu405ts)* were performed at 15°C using the *prg-1(gg540)* mutant. Four hermaphrodites were mated per cross (see Figure S4) and 5 F_1_ heterozygotes were singled for each mated hermaphrodite. Four F_2_s animals homozygous for the naive <u>*dpy-20(e1282) oma-1(zu405ts)*</u> allele were singled for each F_1_ animal and allowed to lay a brood. Four pools of 5 F_3_ animals for each F_2_ were tested for *oma-1(zu405ts)* silencing based on the embryonic arrest assay described above. For the *gfp* paramutation test in Figure 4a, *prg-1(gg531)* mutants were outcrossed with *gfp* expressing animals to remove the *oma-1(zu405ts)* allele, allowing this experiment to be performed at 20°C. Note: because the silenced *oma-1(zu405ts)* and *gfp* alleles are paramutagenic in *prg-1(gg531)* mutants, the wild-type *oma-1* allele in the new strain (YY1256) is also silent, as is the *gfp* allele. Two hermaphrodites were then mated with animals homozygous for the expressed <u>*dpy-10(e128) gfp*</u> allele. Lineages were established from homozygous <u>*dpy-10(e128) gfp*</u> F_2_ animals, which were genotyped for *prg-1(gg531)* once they had laid a brood. Gravid adults were egg prepped every generation to maintain these lineages. *gfp* expression was scored in the F3 and F7 generations using the Plan-Apochromat 20 × /0.8 M27 objective on an Axio Observer.Z1 fluorescent microscope (Zeiss). At least 50 animals were counted for each strain.

### CRISPR/Cas9-mediated genome editing

The CRISPR/Cas9 strategy described previously^4–6^ was used to generate deletions of *rde-3* and *hrde-1*, as well as to introduce the DDH → DAH mutation^7^ in *prg-1*. Guide RNA plasmids and repair template DNA were prepared as described previously^6^. All guide RNAs were designed using the guide RNA selection tool CRISPOR^8^.

### Gene expression quantification using qRT-PCR

Total RNA was extracted from gravid adult animals using TRIzol Reagent (Life Technologies, 15596018). 2ug of total RNA was reverse-transcribed to generate first-strand cDNA using the Superscript III First-Strand Synthesis System (Invitrogen, 18080051) and random hexamers. Note: total RNA was heated with dNTPs and random hexamers to 65°C for 5 mins and immediately chilled on ice before proceeding with remaining cDNA synthesis steps. First-strand cDNA was then treated with RNAse H at 37°C for 20 mins. cDNA was then 1:100 (for *oma-1* and histone RNA quantification) or 1:8 (for all other quantifications) and 2ul was used to quantify gene expression using iTaq Universal SYBR Green Supermix (Bio-Rad) according to manufacturer’s instructions. qRT-PCRs were performed using the CFX Connect machine (Bio-Rad) and semi-skirted PCR plates (Bio-Rad, 2239441). All qRT-PCR data was normalized to quantification of *nos-3*, a germline-expressed gene. qRT-PCR primers are listed in Table 2.

### Taqman-based small RNA quantification

1ug of Trizol-extracted RNA from gravid adult animals (see above) was used for Taqman assays. Small RNAs were reverse transcribed into cDNA using the Taqman MicroRNA Reverse Transcription Kit (Applied Biosystems, 4366596). *oma-1* and *gfp* small RNAs were then quantified by qRT-PCR using TaqMan Universal Master Mix II, no UNG (Applied Biosystems, 4440040) and custom TaqMan small RNA assays from Applied Biosystems (assay IDs: *gfp:* CSLJH0V, *oma-1* probe 1: CSKAJ9W, *oma-1* probe 2: CSLJIF4, *oma-1* probe 3: CSMSGMC, *oma-1* probe 4: CSN1ESK). qRT-PCRs were performed using the CFX Connect machine (Bio-Rad) and semi-skirted PCR plates (Bio-Rad, 2239441).

### pUG RNA detection using pUG PCR

pUG RNAs were detected using pUG PCR, a PCR-based assay described previously^9^. Briefly, total RNA was extracted using TRIzol Reagent (Life Technologies, 15596018). 5ug of total RNA and 1pmol of reverse transcription (RT) oligo (see Table 2) was used to generate first-strand cDNA using the Superscript III First-Strand Synthesis System (Invitrogen, 18080051). Total RNA, dNTPs and RT oligo were mixed and heated to 65°C for 5 mins and immediately chilled on ice before proceeding with remaining cDNA synthesis steps. 1ul of cDNA was used to perform a first round of PCR (20ul volume) with Taq DNA polymerase (New England BioLabs, M0273) for 20-25 cycles and primers listed in Table 2. These PCRs were then diluted 1:100 and 1ul was used to perform a second round of PCR (50ul volume) for 25-30 cycles using primers listed in Table 2. *gsa-1*, which has an 18nt long genomically UG repeat in its 3’UTR, served as a control for all pUG PCR analyses. PCR reactions were then run on 1.5-2% agarose gels. Images were acquired using a ChemiDoc MP Imaging System (Bio-Rad). All image processing was done using Fiji^3^. All pUG PCR reactions were sequenced by cutting out lanes of interest from agarose gels and gel extracting the DNA using QIAquick Gel Extraction Kit (Qiagen, 28706). 3ul of gel extracted PCR product was used for TA cloning with the pGEM-T Easy Vector System (Promega, A1360) according to manufacturer’s instructions. Ligation reactions were incubated overnight at 4°C. Transformations were performed with 5-alpha Competent *E. coli* cells (NEB, C2987H) and plated on LB/ampicillin/IPTG/X-gal plates (according to pGEM-T Easy Vector System manufacturer’s instructions). White colonies were selected on the day next and inoculated in Luria Broth overnight. Liquid cultures were then miniprepped using QIAprep Spin Miniprep Kit (Qiagen, 27106) and plasmid DNA was Sanger sequenced using a universal SP6 primer (5’-CATACGATTTAGGTGACACTATAG-3; Dana-Farber/Harvard Cancer Center DNA Resource Core, Harvard Medical School) or a universal M13 primer (5’-TGTAAAACGACGGCCAGT-3’; Quintarabio, Cambridge, MA).

